# Traction Force Microscopy with DNA FluoroCubes

**DOI:** 10.1101/2024.04.12.589182

**Authors:** Armina Mortazavi, Jianfei Jiang, Philip Laric, Dominic A. Helmerich, Rick Seifert, Svetozar Gavrilovic, Markus Sauer, Benedikt Sabass

## Abstract

Mechanical forces at the cell–substrate interface govern processes from migration to differentiation, yet mapping these forces at high spatial resolution remains challenging. Traction force microscopy (TFM) addresses this by quantifying substrate deformations using fiducial markers, which are conventionally fluorescent beads. Here, we introduce fluorescently labeled DNA nanostructures (FluoroCubes) as alternative fiducials grafted onto polydimethylsiloxane (PDMS) substrates. Co-anchored with RGD peptides, FluoroCubes remain stably tethered, resist internalization, and enable dense, minimally perturbing labeling. This surface-functionalized platform is compatible with TIRF microscopy and leverages tunable biotin–NeutrAvidin chemistry for precise control of fiducial density. Using a modified multi-channel optical flow algorithm, we achieve improved displacement sensitivity and force reconstruction resolution compared to conventional algorithms. FluoroCube-functionalized substrates provide a reproducible, high-resolution method for traction force mapping and offer a versatile foundation for future integration with DNA-based molecular sensors to probe interfacial forces at biointerfaces.

## Introduction

Cell-generated forces are an integral part of many physiological and pathophysiological processes, for example, during embryogenesis, tissue development, wound healing, and immune response^1^. Traction force microscopy (TFM) is a widely adopted method for quantifying these forces due to its versatility, robustness, and ability to capture the dynamic micromechanics of living cells as they interact with their environment^2–4^. Following the seminal work of Harris et al. in the 1980s, who demonstrated that adherent cells could induce wrinkles in thin silicone substrates, Dembo et al. were able to measure spatial maps of cell-generated traction using substrates with a linear elastic response^5,6^. In a conventional TFM experiment, cells are seeded onto an elastic gel containing fluorescent beads as fiducial markers. As the cells interact with the gel substrate, they exert mechanical forces that displace the gel and, with it, the fluorescent beads. Bead displacement is monitored by optical fluorescence microscopy to assess the deformation of the gel relative to its relaxed state. The magnitude and direction of the locally exerted surface stress are then calculated by modeling the mechanical response of the substrate^7^.

With a typical resolution limit of a few micrometers, characterizing traction force patterns in fine detail can be challenging with classical TFM, e.g., if one wishes to study the function of subcellular structures. The accuracy of the reconstructed traction maps depends on the amount of information provided by the measurement of the cell-induced deformations. In particular, the spatial interval at which the deformations are sampled limits the resolution of traction on small-length scales. To visualize traction force patterns on lengths comparable to subcellular structures, displacements must be finely sampled. As a result, several experimental methods have been developed to improve the acquired microscopy data to quantify displacement variations on small length scales. These methods are mainly based on increasing the density of fiducial marker beads and/or using super-resolution microscopy. Examples include the combination of two types of beads with different emission spectra, the use of Stimulated Emission Depletion (STED) super-resolution microscopy, and the use of fluctuation analysis to recover high-resolution images of fluorescent beads with spinning disk confocal microscopy^8–10^. More recently, enhanced tracer density super-resolution microscopy (ETD-srTFM) has been introduced, which improves the conjugation efficiency of the beads on the substrate. The study used structured-illumination microscopy (SIM)-TFM and densities of 15/*μm*^2^ with 100 nm fluorescent beads as markers, corresponding to a sampling interval of 258 *nm* ^11^. Based on numerical simulations performed in the same work, this sampling density would, in principle, allow traction forces to be resolved with a spatial resolution of roughly 0.5 µm, according to the Nyquist theorem. However, this sub-focal adhesion resolution was demonstrated only in simulations and not in experimental traction reconstructions^11^. Thus, a resolution of traction forces on a truly molecular scale, or even on the scale of tens of nanometers, is still far from being achieved with conventional TFM methods that rely on the measurement of substrate deformations with fluorescent beads. Beyond beads, other fiducial probes such as gold nanoparticles (AuNPs) and quantum dots (QDs) have been explored in the field. AuNPs are inherently cell-compatible, as their detection avoids the photodamage and photobleaching associated with fluorescence-based probes. However, their visualization relies on scattering or plasmonic resonance, which requires specialized optical setups, and their functionalization on soft, continuous substrates remains challenging^12–14^. QDs provide high brightness and photostability, but in addition to surface functionalization constraints, they can be cytotoxic at high concentrations and may aggregate or perturb the substrate, limiting dense, stable labeling for traction measurements^15^.

While advances in fiducial markers and imaging methods have enhanced the resolution of TFM, classical bead-based approaches remain fundamentally limited in measuring molecular-scale forces. They report forces averaged over ensembles of roughly 10³ to 10⁵ proteins spanning hundreds of nanometers to several micrometers, making it challenging to detect forces below a few tens of pascals^16^. This spatial averaging means that nanometer-scale force fluctuations at individual adhesion molecules, which are central for mechanotransduction, remain largely inaccessible^17^. To address this limitation, molecular force sensors have been developed^18,19^. These include Förster Resonance Energy Transfer (FRET)-based tension sensors, force-sensitive protein domains, and DNA-based probes capable of reporting piconewton-level forces across individual integrins, cadherins, or other mechanosensitive proteins^20–24^. More recently, super-resolved imaging has enabled visualization of single DNA-based tension sensors within adhesion complexes^25,26^. These tools have provided unparalleled insights into adhesion assembly, catch-bond behavior, and ligand-specific engagement, allowing direct visualization of receptor-level forces and mechanotransduction events that cannot be resolved with classical TFM^27–29^.

Molecular force sensors and classical TFM therefore occupy complementary but largely non-overlapping experimental regimes. TFM excels at mapping whole-cell traction patterns on soft, deformable substrates, whereas molecular sensors quantify forces at single adhesion molecules but are usually restricted to rigid surfaces where substrate deformation is negligible. A promising direction is the emergence of DNA-based mechanoprobes engineered for use within soft, deformable substrates, which could bridge this methodological divide by combining the molecular precision of force sensors with the mechanical relevance of soft-substrate TFM. DNA-based mechanoprobes are particularly attractive for this purpose because their force thresholds, reversibility, and photochemical properties can be precisely programmed. Recent studies have demonstrated hydrogel-integrated molecular tension probes capable of reporting receptor-mediated rigidity sensing, laying the foundation for platforms in which TFM and molecular tension imaging can be performed simultaneously^30,31^. By embedding DNA-based probes within TFM-compatible deformable substrates, such hybrid designs could provide a multi-scale view of cell mechanics, capturing both global traction maps derived from displacement fields and molecular tension signatures at specific receptors that drive emergent cellular behaviors. Motivated by these advances, replacing beads with smaller and more biocompatible fiducial markers, such as DNA-origami-based probs, offers a promising route to enhance both the versatility and spatial resolution of classical TFM approaches.

Alongside experimental substrate engineering and fiducial labeling, which rely on polymer and surface chemistry, soft matter physics, and advanced microscopy, computational analysis is essential for reconstructing cellular traction fields and interpreting mechanobiological data in TFM. To provide context, we briefly introduce the standard procedures used to calculate cellular traction forces from microscopy data. The calculation of cellular tractions is based on pairs of microscopy images containing fiducial markers, such as fluorescent beads, to visualize the substrate deformations in the vicinity of cells with respect to images of the undeformed substrate without cells, which are typically removed by trypsinization. After acquiring high-resolution images of the cell and the fiducial markers, the next step is to quantify the deformations of the substrate. For this purpose, the displacement of fiducials between image pairs is determined using either Particle Tracking Velocimetry (PTV) or Particle Image Velocimetry (PIV). In PTV, the motion of individual fiducial markers is tracked to generate dense displacement fields^32^. However, an accurate localization and assignment of fiducials in the image pairs is necessary for single-particle tracking, which can be challenging for dense fluorescent probes with overlapping point spread functions (PSFs). In contrast, PIV can robustly quantify displacements in an image that is divided into small windows that can contain more than one fiducial. The displacement of each window in one image with respect to another image is obtained by maximizing its cross-correlation with respect to a local, relative shift of the images^33^. Besides PIV and PTV, Optical Flow Tracking (OFT) has recently emerged as a method to quantify the displacements in a TFM experiment with high precision and spatial resolution^34^. Similar to PIV, OFT relies on local image features and does not require precise localization of the individual particles. Object tracking requires interrogation windows, and within each window, OFT requires intensity consistency^35^ and local smoothness^36^. If these conditions are met, dense clusters of fiducial markers can be tracked even if their PSFs overlap strongly^37,38^. However, analogous to PIV, the size of the window affects the smoothness of the resulting displacement field in OFT. Large window sizes result in smooth displacement fields that contain little noise, while small window sizes can capture local details of the displacement field at the cost of being more susceptible to noise. Once a displacement field has been extracted from the data, the spatial map of cell-generated tractions can be computed using various methods. The most widely used methods are based on Green’s functions that relate a displacement field to the application of a point-like surface traction^6,39^. Among these methods, Fourier Transform Traction Cytometry (FTTC) stands out due to its robustness and computational efficiency^8,40^. By calculating traction fields in the Fourier domain, FTTC significantly reduces computational effort, and detrimental noise in the displacement field can be dealt with classical regularization schemes or Bayesian Fourier Transform Traction Cytometry (BFTTC)^41^.

In this work, we move beyond traditional bead-based approaches by employing DNA-based, molecular-sized probes as novel and unconventional fiducial markers for TFM. The DNA FluoroCubes, introduced in Ref.^42^, are approximately 6 nm in size and incorporate multiple fluorophores to enhance photostability. We extend their application to an *in vivo*-like cellular context, demonstrating their stability and suitability for long-term imaging on deformable substrates. The FluoroCubes are immobilized on silicone substrates via biotin–NeutrAvidin interactions, enabling high-quality imaging using total internal reflection fluorescence (TIRF) microscopy^43–45^. In addition to FluoroCubes, fluorescent nanobeads with a complementary emission spectrum are incorporated as a demonstration of feasibility and in combination with the dual-channel setup to achieve high-resolution tracking of substrate displacements. We develop a custom modification of the classical Kanade–Lucas–Tomasi (KLT) optical flow algorithm^35,38^, which analyzes both channels simultaneously and provides validation against conventional fiducials. The performance of these methods is demonstrated using both synthetic and experimental data.

## Results and Discussion

In the following, we present results from the development of FluoroCube-based TFM methods, including experimental procedures and image analysis.

### Imaging FluoroCubes on PDMS substrates

Established materials for cell substrates in TFM experiments are either hydrogels or elastomers, such as polyacrylamide (PAA) or polydimethylsiloxane (PDMS), respectively, which allow for a tunable stiffness roughly between 100 Pa and 50 kPa. For PAA hydrogels used in a standard TFM approach, the fluorescent beads are typically incorporated into the gels by mixing them with the gel constituents prior to polymerization. However, the presence of the beads throughout the gel causes light scattering and diffraction, which can result in a large background fluorescence if the fluorophores are excited outside the focal plane. Additionally, the different axial positions lead to distorted or obscured images of the beads in a wide-field image. Mixing the beads with the PAA gel components can also result in clusters of beads, which appear as bright spots under the microscope and thus prevent the tracking of individual beads. Potentially, the clusters can even introduce unwanted spatial variations in the material parameters that govern the mechanical response of the gel. Overall, the embedding of fiducial markers inside the gels makes accurate data analysis and the interpretation of traction force maps a challenging task. However, fluorescent beads can also be attached only to the gel surfaces, which improves control over their spatial distribution in the image plane and improves data quality^46^.

PDMS substrates provide certain advantages due to their excellent optical transparency, allowing for efficient excitation and detection of fluorophores. Furthermore, the use of a PDMS substrate, with its higher refractive index compared to PAA gel, enables imaging of fluorescent molecules close to the surface using TIRF or a highly inclined illumination mode (HILO, sometimes referred to as “dirty TIRF”)^44,45,47^. In conventional TIRF, an evanescent field is generated at the interface, exciting fluorophores only within approximately 100–200 nm of the surface. In contrast, HILO employs a slightly smaller incidence angle, producing a highly inclined but non-evanescent light sheet that penetrates deeper, up to about 10 µm, which allows visualization of structures below the immediate surface while still reducing background fluorescence. For PDMS substrates made with the Dow DOWSIL™ CY 52-276 Kit and refractive index of 1.403^45,48^, we demonstrated that transitioning from epifluorescence (EPI)-illumination to TIRF significantly enhances the signal-to-background ratio, particularly for FluoroCubes. For FluoroCubes, which have fewer conjugated dyes compared to beads, EPI-illumination fails to produce a distinguishable signal due to the dominance of background noise, rendering these probes nearly undetectable, see Figure 1. Additionally, we compared TIRF imaging with confocal microscopy. While confocal microscopy also offers optical sectioning, its signal-to-background ratio is substantially lower than that of TIRF for these applications. This limitation may arise from a collection of fluorescence signals from a broader axial range, resulting in more out-of-focus light that contributes to the background, see Figure S1. In contrast, optical sectioning with TIRF excites fluorophores exclusively within a narrow region near the substrate surface, effectively eliminating out-of-focus excitation and providing superior imaging quality for smaller probes like FluoroCubes. These findings highlight the critical role of TIRF in enhancing resolution and signal clarity in advanced TFM^49^.

**Figure 1.**
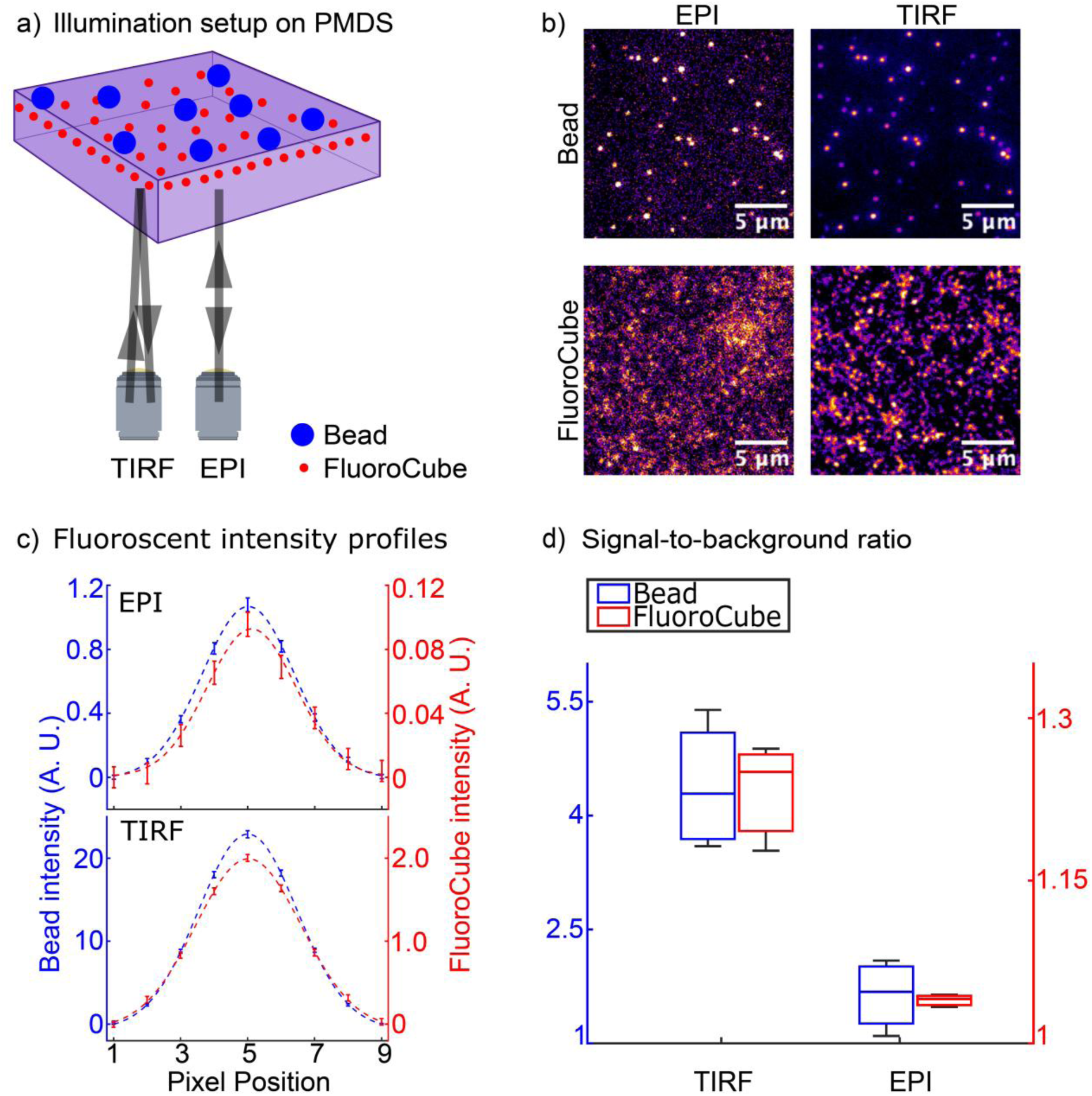
Comparison of image quality in total internal reflection fluorescence (TIRF) and epifluorescence (EPI) microscopy for fluorescent beads and FluoroCubes on polydimethylsiloxane (PDMS) substrates. a) Illustrations of the different illumination setups where 40 nm, red-orange fluorescent carboxylate-modified microspheres (∼0.25 probes·µm^−2^) and FluoroCubes (∼2.5 probes·µm^−2^) are conjugated to the top surface of PDMS gel and imaged with EPI or TIRF illumination mode. b) Enhancement in signal-to-background ratio achieved with TIRF, particularly for FluoroCubes, which exhibit weaker fluorescence signals under EPI-illumination due to their lower dye content. PDMS substrates have an approximate thickness of 10 µm, n = 22 measurements. c) Fluorescence intensity profiles around each probe as an estimate for the point-spread function. Data is averaged over all detected fluorescent probes in the images, 613 beads detected in EPI images, 2245 beads in TIRF, 23549 FluoroCubes in EPI, and 15849 FluoroCubes in TIRF. Dashed lines are fitted Gaussian functions, and data points are average intensities along the central horizontal line of the individual probe with error bars indicating the standard error of the mean. d) Box plots of signal-to-background ratio are obtained from 6 different TIRF and EPI images for FluoroCubes and 8 different TIRF and EPI images for beads.

### Dual labeling of substrates with FluoroCubes and beads

The use of FluoroCubes, which have a size similar to the green fluorescent protein, allows one to increase the density of markers on the substrate, improving the spatial coverage of the traction force maps. However, a FluoroCube with six fluorescent dyes emits far fewer photons than a fluorescent bead containing 100s of dyes, making precise localization more challenging. To validate that FluoroCubes can serve as reliable fiducial markers and improve the localization precision, we additionally labeled the PDMS substrates with fluorescent beads of varying sizes and emission spectra. Figure 2a illustrates the structure of the dual-labeled surface coating. Briefly, the hydrophilicity of the PDMS surfaces is improved through functionalization with amine groups, followed by passivation with biotinylated bovine serum albumin (BSA). This process allows for the conjugation of NeutrAvidin to the substrate, which, in turn, enables high-affinity binding of FluoroCubes with biotin moieties. RGD peptides, as a substitute for larger extracellular matrix proteins, and carboxylate-modified microspheres are then activated with carbodiimide and reacted with amine groups of the substrate to form amide bond derivatives, see Figure 2b^50,51^. Compared to fibronectin coatings, which permit cell-mediated fibrillogenesis that can alter local mechanics and amplify traction readouts, RGD peptides provide a more stable interface for quantifying pure traction forces^52^.

**Figure 2.**
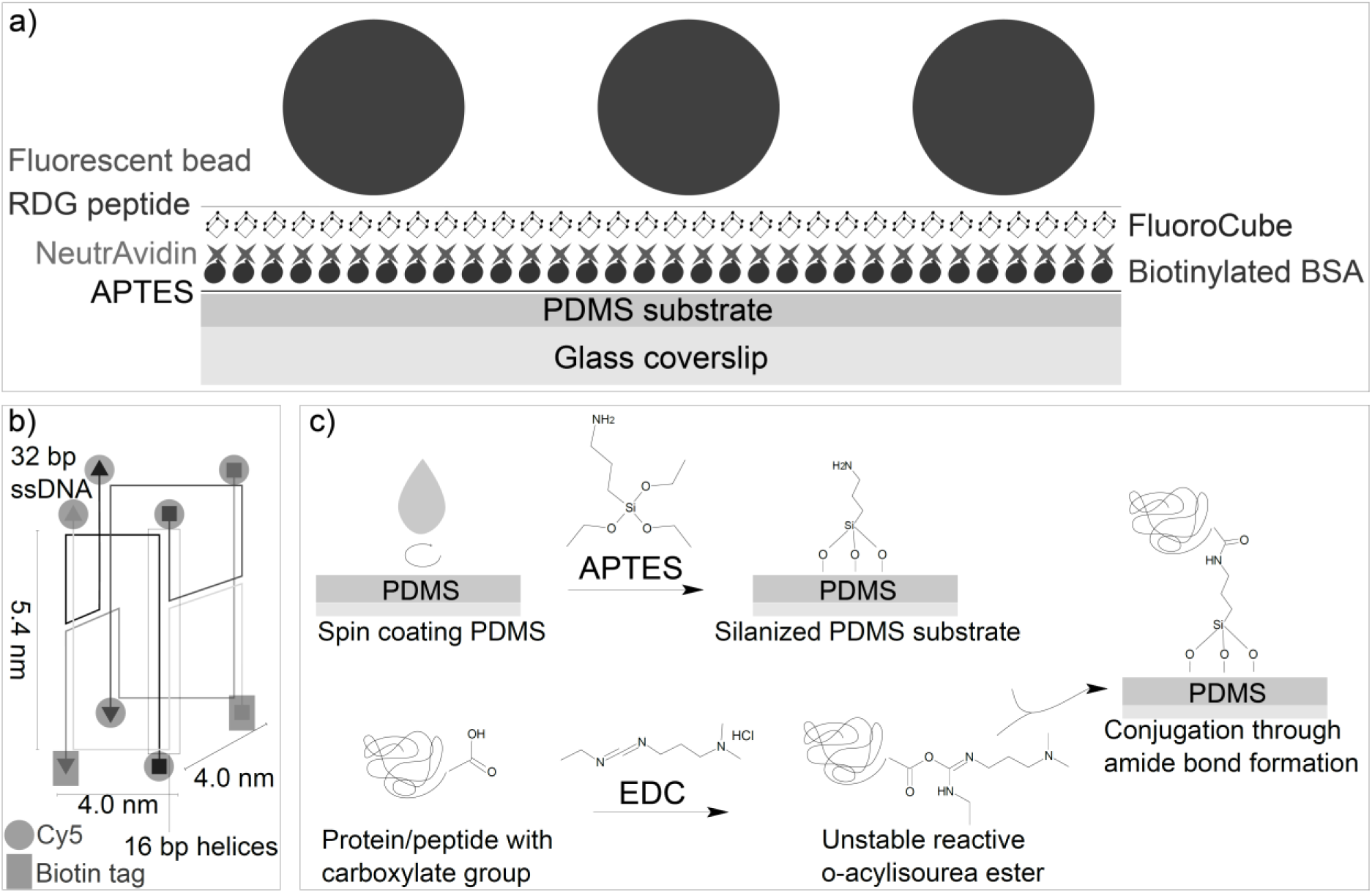
Functionalization of silicone substrates for traction force microscopy with FluoroCubes. a) PDMS substrates (thickness of ∼10 µm) with a refractive index of ∼1.4 are passivated with biotinylated bovine serum albumin (BSA) after functionalization with aminosilane (3-Aminopropyl)triethoxysilane (APTES). As a linker molecule, streptavidin reported in Ref.^42^ was replaced with NeutrAvidin, which forms a complex with FluoroCubes attached to a biotin tag at one site and biotinylated BSA at another site. 40 nm carboxylated beads are bound to the substrate, and RGD peptides are grafted to the substrate for cell adhesion. b) FluoroCubes comprise six Cy5 dyes and two biotin molecules, all conjugated to 32 bp ssDNA that is folded subsequently. c) Conjugation of beads and peptides starts with activation of carboxylate by 1-ethyl-3-(3-dimethylaminopropyl) carbodiimide (EDC). EDC-mediated conjugation leads to the formation of a highly reactive *O*-acylisourea intermediate that forms an amide bond with amine groups on the surface of the PDMS substrate.

The setup is based on a newly developed biotin–NeutrAvidin functionalization strategy for PDMS substrates that enables combined TIRF imaging and TFM with DNA-based probes. Although we demonstrate the feasibility using FluoroCubes, fluorescent beads, and a dual-fiducial configuration, the approach is not restricted to this specific set of probes. Any biotinylated or carboxylated fiducial marker, including alternative DNA-based nanostructures, could be employed. This provides a versatile platform that can support future multi-color and high-density TFM applications.

### Modified optical flow tracking for improved traction-force resolution with dual-labeled samples

For the inference of a displacement field, we compare images of beads and FluoroCubes in the presence of a cell with images of the same region after removal of the cell by trypsinization. A versatile Kanada-Lucas-Tomasi (KLT) optical flow tracking algorithm is used to analyze the images (see Methods). However, the standard KLT algorithm can only track displacements from pairs of images recorded in one fluorescence channel, e.g., only from images of the beads.

Therefore, we performed tests with synthetic data to find out how to optimally extract coherent deformation fields from pairs of images recorded in the two channels corresponding to FluoroCubes and beads. The synthetic data consisted of high-resolution displacement fields (Figure 3ai) and traction fields (Figure 3aii), where the displacement fields were generated by a pre-defined traction field and based on an area density of markers of around 5 *μm*^−2^ to replicate the images obtained from our experimental setup. We analyzed the displacement fields obtained from different OFT schemes and also compared the corresponding traction fields as illustrated in Figure 3. Note that post-processing strategies could, in principle, be used to exclude tracking errors from the final data. However, this was not done here, as our goal was to directly evaluate the performance of the modified KLT algorithm itself.

**Figure 3.**
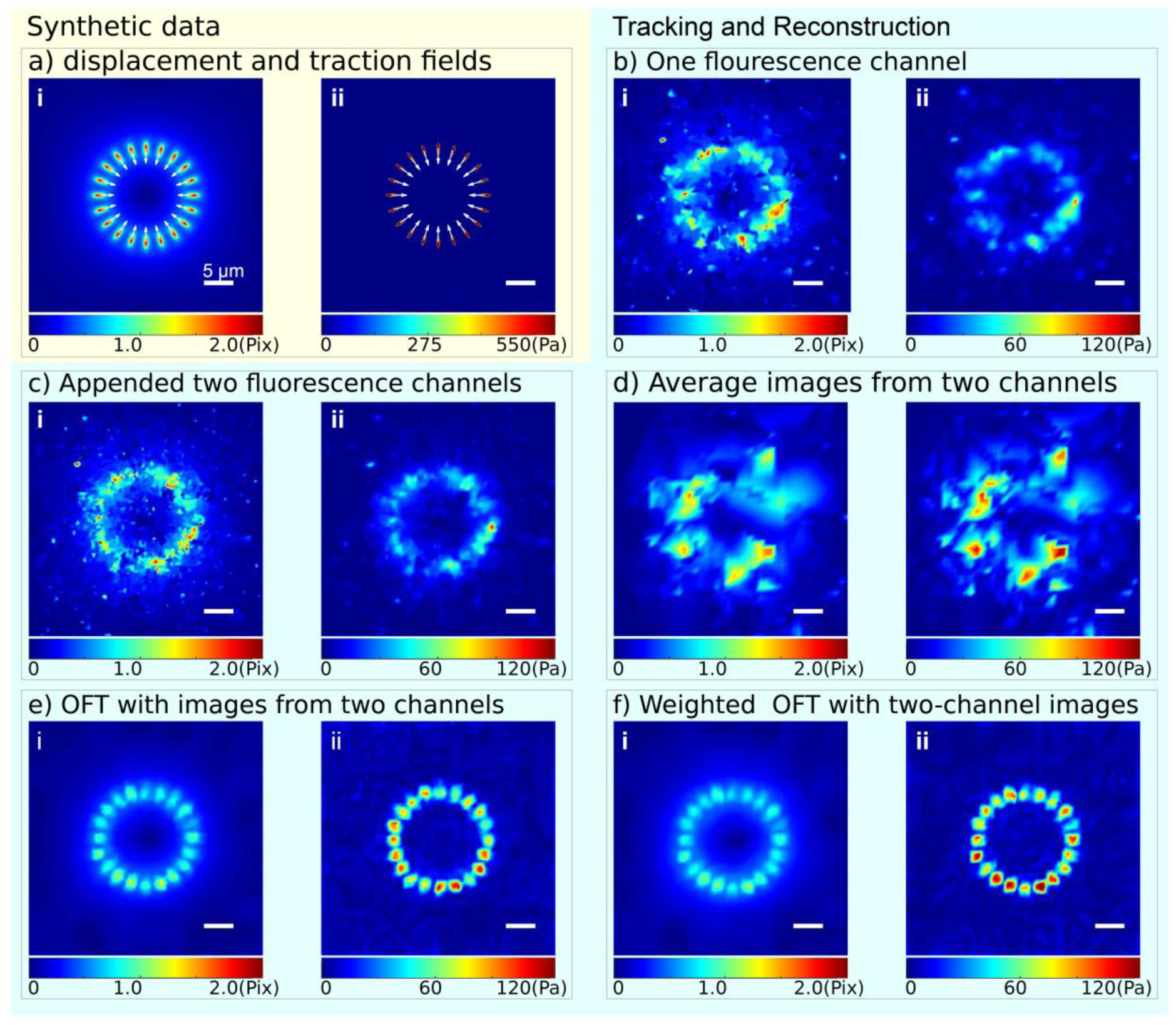
Optimization of TFM resolution by combining information from two fluorescence channels requires a modified KLT tracking algorithm. The two channels correspond to images of FluoroCubes and beads, respectively. a) Synthetic displacement field (i) generated from a pre-defined traction pattern (ii); b) Displacement field resulting from OFT in only one of the two channels (i) and the traction field reconstructed from this data (ii); c) Displacements resulting from independent OFT in both channels and subsequent appending of both data sets (i) and tractions reconstructed from this data (ii). Note the large noise resulting from independent tracking of both channels; d) Displacements resulting from OFT in images that were created by the superposition of the fluorescence signals from both channels (i) and reconstructed tractions (ii). Averaging fluorescence signals in the images degrades traction resolution since trackable features in individual images are covered and distorted; e) Displacement field extracted from both channels with modified KLT tracking routine (i) and resulting traction field (ii). Concatenation of the two datasets in the linear equations used for KLT improves tracking; f) Displacements extracted from both channels using a modified KLT tracking routine with cross-correlation-based weighting factors (i) and its corresponding traction field (ii). Color bars indicate the magnitudes of displacement and traction fields. Scale bar: 5 µm. All traction fields are reconstructed by the Bayesian Fourier transform traction cytometry (BFTTC).

Use of either of the two channels, representing FluoroCubes or beads, allows one to reconstruct traction forces at rather low resolution, where individual spots of traction are not clearly discernible (see Figure 3b). As a first attempt to combine information from both pairs of images, we tracked the displacements in both channels independently and appended the results, which in principle provides a denser field. However, this approach hardly improved the spatial resolution of traction forces since tracking errors in the two independent channels result in strong local misalignment and magnitude mismatch of the displacements from the two channels (see Figure 3c). Similarly, averaging the images from two fluorescence channels did not improve the resolution because meaningless cross-channel correlations appear in the objective function and because image superposition smooths out intensity variations that are required for tracking, see Figure 3d. However, an extension of the optical flow tracking algorithm to optimize coherent motion in both channels eliminated this issue, resulting in reliable displacement and traction fields with high resolution (see Figure 3e). Furthermore, a cross-correlation-based weighting scheme was applied to take into account different levels of uncertainty in the two sets of data, which further enhanced the reliability and resolution of the traction force fields (see Figure 3f). Since the modified KLT algorithm produces displacement data with minimal tracking noise, the reconstruction of traction forces necessitates only a small amount of regularization (Table S1).

Comparison of the resulting estimated displacement vectors with the ground truth revealed a consistent underestimation of the magnitude. This underestimation is also common for PIV-based methods and can be attributed here to the smoothness assumptions intrinsic to the KLT optical flow method. Additionally, a further systematic underestimation of the traction magnitudes is induced in the traction reconstruction (Figure S2) by regularization-based FTTC methods^8,40,41^.

Under experimental conditions, photobleaching and intensity fluctuations may compromise the displacement tracking. To evaluate the robustness of our modified KLT algorithm under such conditions, we generated synthetic image pairs with randomized intensity and introduced a 5% loss of trackable fiducials in the deformed images to simulate photobleaching. When processing this data, the modified KLT method outperformed the conventional single-channel KLT, successfully reconstructing the traction fields even under degraded imaging conditions (Figure S3). Thus, the dual-channel formulation of our algorithm improves its reliability and performance in the presence of noise.

### Photostability and suitability of FluoroCubes for TFM experiments

While the photophysical properties of FluoroCubes have been extensively characterized *in vitro*^42,53,54^, their stability and reliability have not yet been characterized in a cellular context on deformable PDMS substrates. To evaluate the suitability of FluoroCubes for long-term TFM experiments, we quantitatively assessed their photostability and the stability of their surface attachment in comparison with conventional fluorescent beads. In addition, we demonstrate that FluoroCubes offer the advantage of minimal internalization by cells, providing more consistent fiducial labeling in conditions where bead internalization occurs.

Under continuous TIRF illumination, the photostability of FluoroCubes was directly compared to that of fluorescent beads excited in the same channel under identical conditions. Fluorescence intensity was computed as the average estimated photon number per localization over detected probes in each frame and normalized to the corresponding maximum photon number. As shown in Figure 4a, both probe types exhibit a rapid initial decrease in normalized intensity followed by a markedly slower decay at later times. Neither FluoroCubes nor beads reach half of their maximum normalized intensity within the observation window of 2,000 seconds, demonstrating reasonable photostability of both probes under these conditions in the optimized buffer. The observed decay profiles deviate from a single-exponential behavior, suggesting that multiple photophysical processes contribute to the intensity evolution. Note that while the beads remained slightly brighter after prolonged exposure to laser light, the employed continuous illumination for 2,000 seconds represents a rather extreme stress test compared with typical time-lapse TFM experiments, during which imaging is performed intermittently. In standard TFM, photobleaching is relevant insofar as it affects fiducial trackability and force reconstruction. However, under our experimental conditions, it did not impose noticeable practical limitations on temporal or spatial resolution.

**Figure 4.**
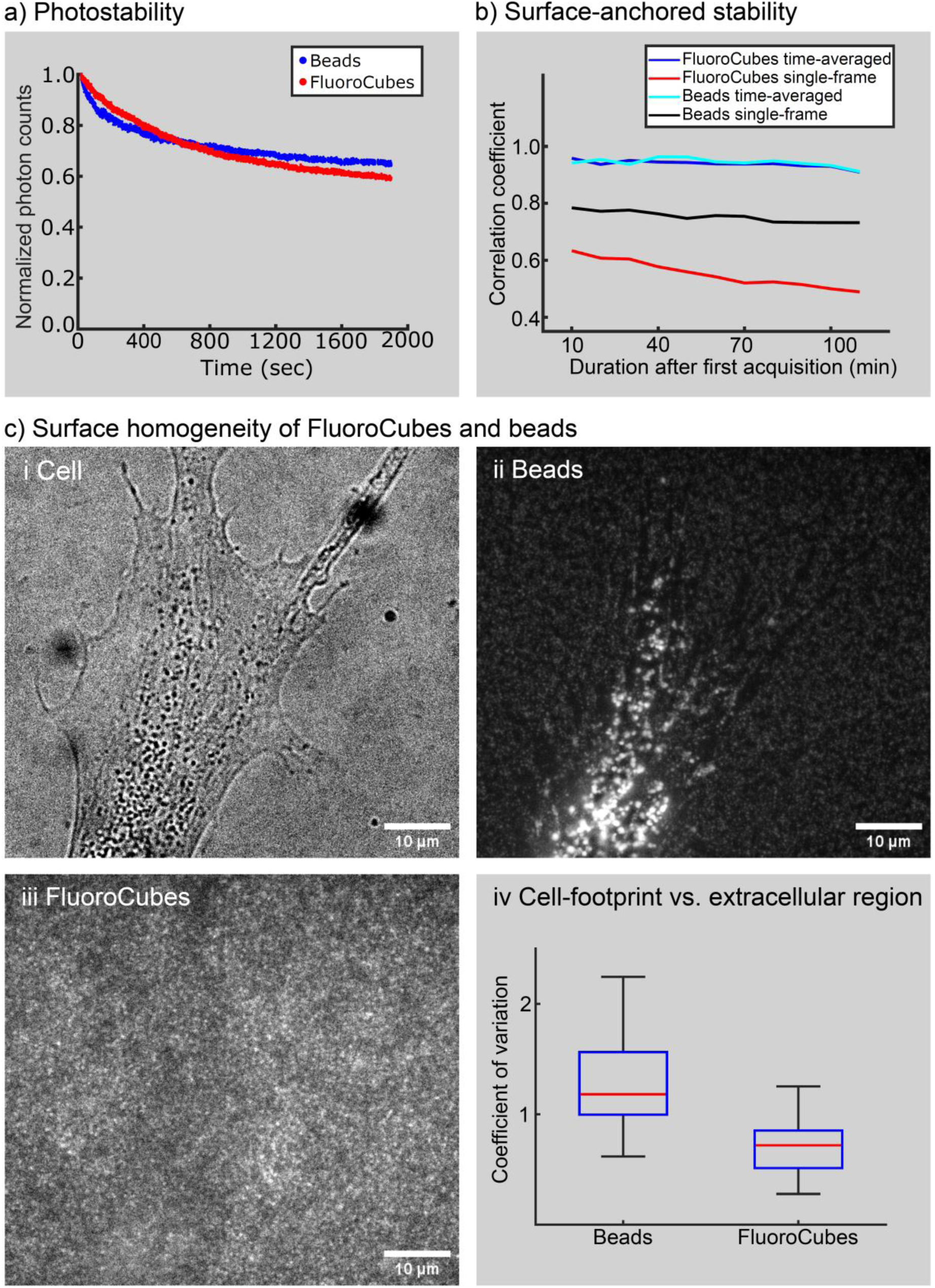
Characterization of the reliability of FluoroCubes as fiducial markers for TFM. a) Normalized fluorescence intensity of FluoroCubes and 40 nm fluorescent beads measured under continuous TIRF illumination using the same excitation channel and identical laser power. b) Maximum cross-correlation between each image and the initial acquisition over 100 min, computed for both time-averaged and single frames. Cross-correlation values close to 1 indicate high surface stability and minimal temporal changes in the fiducial layer. c) Label-homogeneity beneath cells. (i) Bright-field image of a representative cell. (ii) Bead fluorescence. (iii) FluoroCube fluorescence. (iv) Quantification of surface homogeneity using the coefficient of variation (CV) of the mean intensity over non-overlapping 32×32-pixel windows, n = 21 cells. Lower variability reflects a more uniform and stable distribution of labels beneath cells. The CV in areas covered by a cell was normalized by the CV in exterior regions. A local accumulation of detached fluorescent probes results in a normalized CV larger than unity.

To further assess the reliability of the probes in long-term experiments, we imaged the same field of view every 10 minutes for 2 hours, followed by re-imaging after 16 hours of incubation in complete DMEM. For these time-lapse experiments, we quantified the maximum normalized cross-correlation coefficient (NCC) for FluoroCubes and 40 nm red-orange beads using time-averaged and individual frames from independent experiments. The maximum NCC represents the degree of similarity between a given image and a reference template image. In our analysis, the first acquisition was used as the template, and the maximum NCC was computed for all subsequent acquisitions. As shown in Figure 4b, the NCC gradually decreased for both FluoroCubes and beads over time (>100 min) when individual frames were used, reflecting the effects of intensity fluctuations and photobleaching. In contrast, both channels remained strongly correlated when time-averaged frames were used, indicating that averaging a few rapidly acquired images effectively suppresses noise arising from temporal intensity variations. Figure S4 further shows that different averaging intervals yield highly similar images, confirming that the improvement in image quality is rather insensitive to the specific choice of time-averaging. Notably, the maximum NCC for FluoroCubes remained stable over the entire imaging period of 2 h and we measured a maximum NCC of 0.8781 after 16 h when time-averaging was used. This result suggests that FluoroCubes remained firmly attached and consistently trackable with negligible positional drift or loss of signal. Furthermore, surface density measurements performed after 3 days and one week in DMEM revealed minimal fluorescence loss. The biotin–NeutrAvidin coating and conjugated FluoroCubes also remained stable for several weeks when stored in PBS at 4 °C.

Since FluoroCubes contain only a few dye molecules, they naturally exhibit stochastic intensity fluctuations. With beads, such fluctuations are averaged out by simultaneous emissions occurring among the large number of dye molecules in one probe. Given the observed blinking behavior of FluoroCubes, we employed direct stochastic optical reconstruction microscopy (*d*STORM) to precisely localize and track individual FluoroCubes^55^. However, for densely packed FluoroCubes, it was impossible to reliably assign each label in the undeformed configuration to a corresponding label in the deformed configuration. We attribute this limitation primarily to fluorophore crosstalk, possibly via FRET^56^, which modulates dye brightness. Consequently, super-resolution tracking of fluorophores was feasible only for sparsely distributed probes, and dSTORM offered little advantage over diffraction-limited imaging in terms of spatial traction-force resolution. Video S1 illustrates an example of low-density probes that can be tracked individually. Various biocompatible STORM buffers were tested, but did not improve the blinking behavior of Cy5 dyes associated with FluoroCubes sufficiently to reliably use *d*STORM for TFM. In addition, the high-intensity laser and long acquisition time of *d*STORM can affect the cells’ morphology, which in turn changes the traction patterns^57^.

For high-resolution TFM, achieving a dense and uniform distribution of fluorescent probes is essential. However, the integrity of the surface labeling can be compromised beneath cells due to the removal or internalization of probes, particularly for highly migratory cells or cells with particularly high mechanical activity. Such perturbations confound TFM analysis, as internalization not only reduces the number of trackable fiducials but frequently creates regions beneath the cells where tracking is impossible due to spurious fluorescence signals. In our experiments, the extent of bead internalization varied markedly between individual cells and across cultures, influenced by factors such as cell type, incubation duration, and passage number. To determine whether FluoroCubes, like beads, are subject to detachment or uptake by cells, we compared the spatial homogeneity of their fluorescence signals beneath the cells to that of outside the cells. Specifically, we quantified the normalized coefficient of variation (CV) for both beads and FluoroCubes (Figure 4c, see Methods). The normalized CV in bead images exceeds 1 (median ∼1.18), indicating that local bead intensities vary more in cells than in the surrounding substrate. This increased variability likely reflects internalization or repositioning of the beads by the cell. In contrast, FluoroCube images show the opposite trend, with a median CV of approximately 0.7, indicating that intensity variations are lower within the cell footprint compared to extracellular regions. The deviation from a normalized CV of 1 for FluoroCubes arises from slight defocusing under the cell body, where vertical forces deform the substrate and shift some FluoroCubes marginally out of the optimal focal plane^58^, as also visible in Figure 4c) iii. For additional verification in a subsample, we manually quantified the number density of FluoroCubes beneath the cells and in the surrounding region. The densities were nearly identical (1.93 µm⁻² beneath the cells and 1.89 µm⁻² outside), consistent with the absence of FluoroCube uptake. In contrast, bead densities showed greater variation, with 1.37 µm⁻² beneath the cells and 2.02 µm⁻² outside the cells, in line with their tendency to be internalized by cells. This analysis was exclusively performed on cells in which beads were confirmed by confocal z-stacks to be internalized or repositioned by the cell. In cases where the beads’ signals did not accumulate inside the cell, FluoroCubes also did not appear in intracellular z-stacks. These results suggest that, unlike beads, FluoroCubes remain stably immobilized, are less susceptible to cell-induced detachments, and remain extracellular, thereby providing a more reliable and uniform fiducial layer for TFM experiments; whereas DNA nanostructures that are internalized are generally susceptible to nuclease-mediated degradation^59^. Consequently, issues associated with internalization of surface markers and tracking errors from intracellular fluorescence signal do not arise with FluoroCubes^60^.

Although these results demonstrate robust attachment and photostability of DNA-based FluoroCubes under the tested conditions, we note for completeness that labeling integrity in nuclease-rich environments can be further enhanced through additional protection strategies^61^. Short-term protection can be achieved using sacrificial “decoy” DNA, where excess unmodified duplex DNA acts as a competitive substrate for nucleases, protecting functional constructs from degradation^62^. Long-term stability can be enhanced through chemical backbone modifications such as phosphorothioate (PS) linkages, L-DNA analogues, or peptide nucleic acids (PNA), which increase resistance to enzymatic cleavage while maintaining binding functionality^63,31,64^. Incorporating one or both strategies provides a flexible framework to extend the durability of DNA-based constructs in harsh, nuclease-rich environments without compromising surface attachment or photostability.

Collectively, these results demonstrate that FluoroCubes are highly photostable, can be bound stably to a surface, and are not taken up by cells, enabling reliable and long-term traction force mapping.

### High-resolution TFM on substrates labeled with FluoroCubes and beads

Next, we demonstrate that the modified setup can also be employed for high-resolution TFM experiments. For experiments with cells, coverslips were spin-coated with PDMS elastomer, and the sample was subsequently affixed to sticky chambers containing 8 or 18 wells. Inspired by methods proposed in Ref.^65^, this setup enables efficient preparation of sample replicates that can be imaged separately or during a single session. The densities of both types of fluorescent probes were carefully adjusted to be as high as possible, while still allowing the distinction of individual fiducial markers with a diffraction-limited microscope for subsequent motion tracking. The resulting number densities, which determine the spatial resolution of traction forces, can further be varied depending on the image acquisition and reconstruction technique (Figure S5)^9,10,49^. The stability of the fluorescence signal from our probes is determined by experimental conditions such as laser power, probe density, local FRET interactions, and buffer composition (e.g., pH or oxygen concentration in the glucose scavenger system). To reduce photobleaching and blinking of the FluoroCubes, Trolox, a triplet-state quencher, was added to the imaging buffer, which additionally contained high-glucose DMEM supplemented with glucose oxidase and catalase^66,67^. Imaging the FluoroCubes in this optimized buffer, which couples a glucose scavenger system with Trolox, substantially reduced photobleaching and blinking (see Video S2; compare with Video S1), both of which otherwise pose challenges for long-term fluorophore imaging. Under these conditions, the fluorescence signal of the FluoroCubes became nearly as stable as that of fluorescent beads (see Video S3), exhibiting only minor intensity fluctuations over time. We also tested an alternative enzymatic oxygen-scavenging system consisting of protocatechuic acid (PCA) and protocatechuate-3,4-dioxygenase (PCD) together with Trolox. The PCA/PCD system is often favored for its long-term stability and reduced buffer acidification during imaging, which can be advantageous for extended time-lapse acquisition^68^. However, in our hands, this system did not provide a consistent improvement in FluoroCube photostability under the tested conditions. Nevertheless, the optimized imaging buffers minimized the fluctuations and kept the localization and tracking performance reliable throughout acquisition. Thus, despite their substantially lower photon budget compared with beads, FluoroCubes can be detected robustly in suitably chosen conditions. To further improve the signal-to-noise ratio for both FluoroCubes and beads at high label density, rapidly acquired successive frames were averaged prior to using the images for tracking.

To quantitatively evaluate the consistency of displacement fields obtained with different surface labelings, we first performed experiments in which the microscope stage was shifted laterally by approximately one micrometer. The displacements inferred for these experiments from images of beads and FluoroCubes were in excellent agreement, as illustrated by the straight line with a slope close to unity that results from plotting the two values against each other, see Figure 5b). The same analysis was subsequently used to compare different algorithms for the measurement of displacements, including cross-correlation PIV, the conventional KLT algorithm and the modified KLT algorithm. All methods produced highly consistent results (Figure S6).

**Figure 5.**
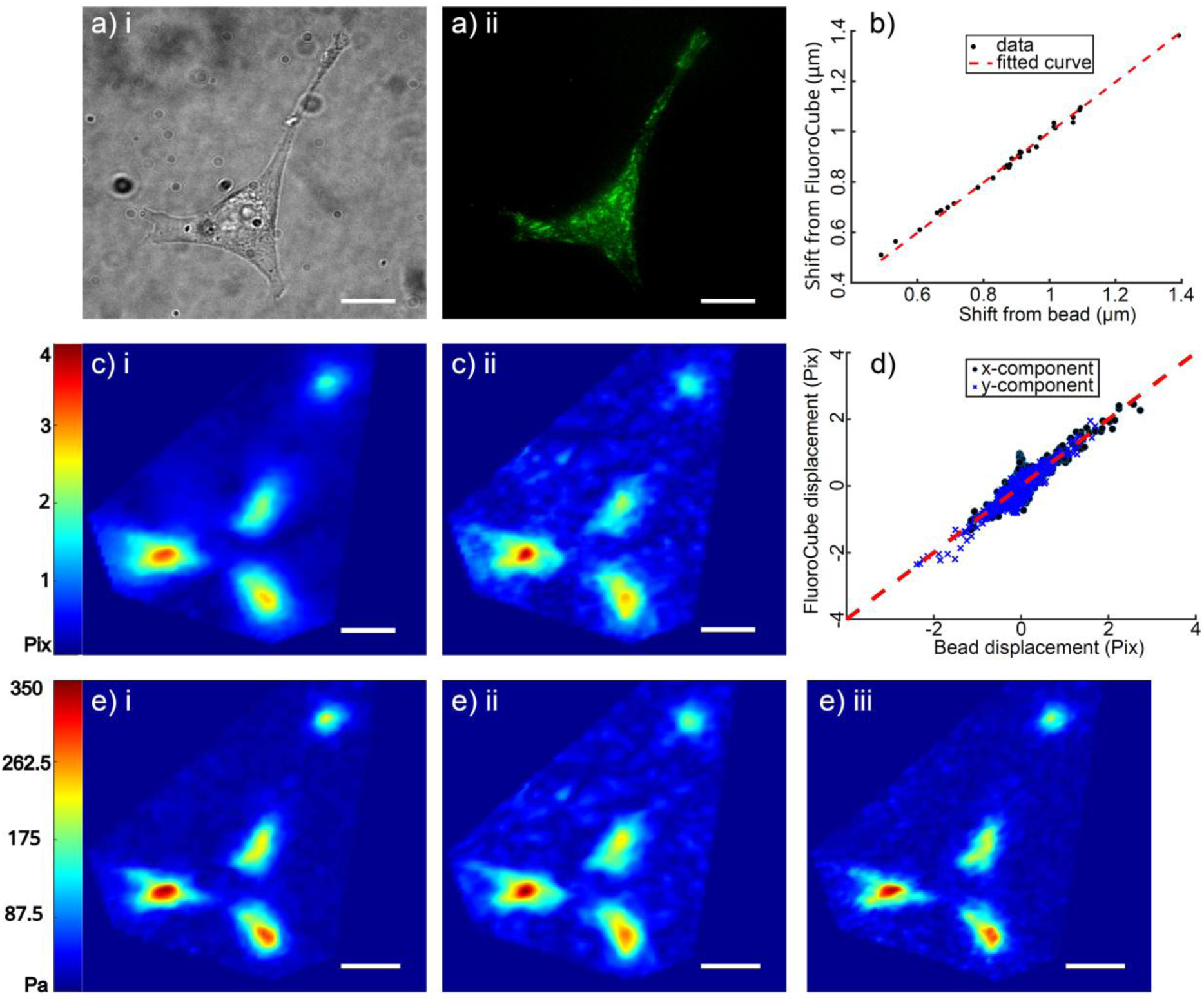
Demonstration of TFM with FluoroCubes and fluorescent beads as fiducial markers. a) (i) and (ii) Bright field image of a murine kidney fibroblast with GFP-labeled kindlin-2 (Kind^Ko+K2GFP^ cell line) and the corresponding fluorescent image; b) Displacement correlation between the bead and FluoroCube channel resulting from the stage shift in the x direction. The black dots represent the shifts measured by the conventional KLT method from the FluoroCube channel as a function of the corresponding shifts in the bead channel. The fitted curve has a slope of 0.9975 with *R*^2^ = 0.9940;; c)-(i) and (ii) Displacement fields measured from the bead and FluoroCube channel, respectively; d) Displacement correlation between the two channels in the experimental images; The micrometer-to-pixel factor is 0.103 *μ*m/pix; e)-(i) Tractions calculated from the displacement in c)-(i) with BFTTC; e)-(ii) Tractions calculated from the displacement in c)-(ii); e)-(iii) Tractions calculated from dual-channel optical flow tracking results (see Figure S7). All scale bars are 12.5 μm.

Next, we evaluated our approach by performing TFM measurements for murine kidney fibroblast cells stably expressing GFP-labeled kindlin-2 (Kind^Ko+K2GFP^), as shown in Figure 5a. Displacement fields measurements using conventional KLT method with a 30-pixel tracking window for the two channels (Figure 5c) demonstrated qualitative agreement, which was further supported by quantitative displacement correlation analysis (Figure 5d). While reducing the tracking window size can enhance the resolution of the displacement field, it may potentially also introduce tracking errors due to the high density of the FluoroCubes (Figure S7a-b). Our modified KLT algorithm mitigates these limitations by simultaneously analyzing both channels, thereby enabling accurate displacement estimation in densely labeled regions (Figure S7c).

Traction fields were reconstructed using BFTTC with a Green’s function that accounts for finite substrate thickness, yielding smooth, high-resolution maps of cellular forces, see Figure 5e^69–72^. The reliability of the traction reconstruction was assessed by examining the spatial co-localization between the calculated traction forces and the fluorescence signal of labeled focal-adhesion proteins in Kind^Ko+K2GFP^ cells on ∼12 kPa PDMS substrates (Figure S8). Kindlins are core focal-adhesion proteins that, together with talin, bind the cytoplasmic tails of β-integrins and link the integrin tails to the contractile F-actin cytoskeleton^73,74^. Therefore, traction forces transmitted by integrins are expected to localize near accumulations of kindlin. As shown in Figure 5e, traction forces calculated from each individual channel were localized near focal adhesion sites, while the combined analysis of both channels provided a more detailed reconstruction of the traction fields due to the increased number of trackable data points. Ecto-tag β1 integrins expressing GFP in murine kidney fibroblasts were also used in these experiments to confirm the colocalization with the focal adhesion sites (Figure S10-12)^75^.

While our experiments combined Cy5-labeled FluoroCubes with red-orange fluorescent beads to demonstrate the performance of the dual-channel optical flow tracking algorithm, FluoroCubes are primarily intended as an alternative rather than a complementary fiducial marker (Figure S13). The multi-channel algorithm itself is independent of the marker type, meaning that two spectrally distinct FluoroCube populations could, in principle, provide equivalent or even superior displacement resolution without the limitations associated with beads, such as their larger size, variable embedding depth, or cellular internalization. This flexibility highlights the potential of FluoroCubes as fully self-contained fiducial systems for high-resolution, live-cell TFM. Future work will explore multi-color FluoroCubes configurations to further enhance spatial resolution and enable multiplexed mechanical measurements on continuous soft substrates.

## Conclusion

Classical TFM methods based on the measurement of substrate deformation are well-established and readily available in many laboratories. They mostly require standard laboratory equipment and are quite versatile^76^. For example, a wide range of sample sizes can be studied, from very small cells to large multicellular systems^77^. Sophisticated analysis methods make it possible to study cellular forces in three-dimensional geometries or on complex topographies^7^, as well as in the presence of external perturbations^78^. Classical TFM produces vectorial traction maps and a wide range of force magnitudes can be detected, starting from a few pN measured for bacterial appendages on soft substrates to forces of up to 600 nN for platelet adhesions that are important for blood clot formation^79–81^. Finally, a relatively high sample throughput can be achieved by leveraging microfabrication techniques to create regular fiducial marker patterns. This approach allows substrate deformations to be quantified directly, without the need to acquire a separate undeformed reference image^82,83^.

Two potential drawbacks of classical TFM are its limited spatial resolution and the requirement for substrate engineering and surface functionalization, which restrict the arbitrary tailoring of the mechanical and chemical properties of the substrate. We refined the classical TFM workflow both experimentally and computationally by introducing DNA-based probes as a novel strategy for nanoscale labeling, together with an improved KLT algorithm for measuring substrate deformations. Conventional fluorescent beads were shown to be replaceable by DNA-based FluoroCubes, multichromophoric nanostructures containing only a few dye molecules each, tethered to biotin–NeutrAvidin functionalized PDMS substrates with negligible pore size. A key advantage of our biotin–NeutrAvidin functionalization strategy is its modularity, allowing straightforward incorporation of alternative DNA-origami or multichromophoric probes and tunable labeling density, giving precise control over surface properties. FluoroCubes were chosen as fiducials due to their photostability and brightness, as demonstrated in *in vitro* studies, which support high-precision particle tracking^42,53,54^. In this work, we extend their use to an *in-vivo*-like cellular environment, leveraging their small size, monovalent attachment, and favorable photophysical properties for TIRF imaging and TFM. To establish that FluoroCubes can be used to robustly reconstruct traction forces, we performed multiple experiments with both beads and FluoroCubes as labels and showed that both approaches yield comparable results.

Simultaneous tracking of fluorescent probes with different emission spectra can improve the spatial resolution of TFM. Having already performed experiments with two kinds of labels, we sought to combine FluoroCube tracking with bead-based tracking. To optimize displacement extraction, we refined the KLT optical-flow algorithm, which now incorporates a dual-channel iterative weighting scheme. The dual-channel optical-flow algorithm can be used with any two spectrally distinct probes, including beads, FluoroCubes, or other fiducial labels. The resulting improvement in displacement sensitivity and force reconstruction is particularly beneficial when working with dense nanoscale fiducials, where very small displacements must be resolved with high fidelity. In our experiments, the resulting high-resolution traction-force maps revealed a pronounced spatial correlation between traction magnitude and the localization of kindlin-2 or β1 integrin at focal adhesions.

Table 1 compares the relative advantages of DNA FluoroCubes and beads for TFM. Each approach has distinct strengths and limitations: FluoroCubes provide a minimal perturbation of the surface topography and elastic response, as well as nanoscale engineering flexibility and modular design, but currently face limitations in brightness and tracking precision compared to beads. An additional benefit is that FluoroCubes remain stably anchored to the substrate, in contrast to conventional fluorescent beads, which are prone to cellular uptake. While beads exhibit higher intrinsic photostability, FluoroCubes’ photostability is sufficient for standard TFM acquisition schemes and does not limit force reconstruction. Beads, although bright and easy to track, potentially perturb the local surface properties and cannot be arranged at densities corresponding to inter-particle spacings below ∼100 nm. When paired with super-resolution microscopy, DNA-based probes offer a route to molecular-scale traction mapping approaching tens of nanometers.

**Table 1.** Comparison of DNA FluoroCubes and fluorescent beads as fiducial markers for TFM.

| Feature / Property | DNA FluoroCubes | Fluorescent Beads |
| --- | --- | --- |
| Size | (+) Very small, ~6 nm<sup>a</sup> | (-) Large, 40–200 nm<sup>b</sup> |
| Brightness | (-) Brighter than individual dye molecules. Less bright than beads<sup>c</sup> | (+) Very bright<sup>c</sup> |
| High-density labeling | (+) Dense labeling possible<sup>d</sup> | (-) Dense labeling possible but bead interactions can affect mechanics<sup>e</sup> |
| Probe homogeneity | (+) DNA self-assembly ensures uniform stoichiometry and size<sup>f</sup> | (-) size and brightness may vary from batch to batch<sup>b</sup> |
| Cellular uptake | (+) Internalization not observed<sup>g</sup> | (-) Prone to cellular uptake<sup>g</sup> |
| Customizability | (+) Geometry, size, brightness, and modularity can be tailored<sup>h</sup> | (-) Fixed geometry; limited spectral options and surface modifications |
| Single-molecule imaging | (+) Compatible with single-molecule fluorescence applications in principle<sup>i</sup> | (-) Not compatible due to the ensemble fluorescence of multiple fluorophores |
| Probe accessibility | (-) Not widely available | (+) Established commercial sources |
| Future potential for nanoscale engineering | (+) Further sensing capabilities possible within probes (pH, ions, molecular force sensors)<sup>i</sup> | (-) Not easily feasible within beads |

<sup>a</sup> Size defined by DNA nanostructure design and prior characterization<sup>42</sup>.
<sup>b</sup> Manufacturer specifications and literature values commonly used for fluorescent beads.
<sup>c</sup> Photobleaching behavior and relative brightness quantified in Figure 4 and Ref. 42.
<sup>d</sup> Demonstrated by tunable surface density and uniform coverage on deformable substrates (Figure 2).
<sup>e</sup> Represented in Ref. 11.
<sup>f</sup> DNA self-assembly and modular probe design described in prior work<sup>42</sup>.
<sup>g</sup> Cellular uptake assessed experimentally (Figure 4c).
<sup>h</sup> Designed and characterized previously<sup>42</sup>.
<sup>i</sup> Prospective or principle-based capability supported by prior studies; not directly demonstrated in this work.

Recent high-resolution force-measurement strategies rely on molecular force sensors directly linked to adhesion ligands on rigid substrates. Beyond their role as fiducials for TFM, DNA nanostructures provide opportunities to integrate molecular force-sensing motifs directly into deformable substrates^84^. Unlike beads, DNA constructs can be engineered to carry multiple fluorophores, mechanical springs, force-sensitive motifs, or environmental sensors^85^. Their modular architecture enables dual-function designs that combine high-resolution displacement tracking with receptor-level mechanotransduction readouts. Such hybrid substrates could connect piconewton-scale molecular events to micron-scale traction patterns, providing multi-scale insights into how cells transmit and interpret mechanical cues with minimal perturbation of the cellular microenvironment.

Future work will explore brighter and more photostable dyes such as Cy3B or Alexa Fluor 647, which are compatible with super-resolution imaging, as well as nuclease-resistant architectures and multicolor FluoroCube pairs that could fully replace beads and eliminate limitations related to bead size, internalization, and labeling density^61,62^. Together with optimized dyes, imaging conditions, and protective DNA modifications, DNA-based fiducials provide a versatile platform for high-precision, long-term live-cell traction force microscopy. Overall, the established methods provide a toolbox for integrating TFM with molecular-scale fluorescent probes, paving the way for advanced setups that combine force measurement with molecular sensors to study diverse cellular signals on surfaces.

## Materials and Methods

### Cell culture

Human foreskin fibroblasts (ATCC; SCRC-1041) and Kind^Ko+K2GFP^ cells (Kind^Ko^ immortalized murine kidney fibroblast reconstituted with green fluorescent protein-tagged kindlin-2 expression plasmids, generous gift from Prof. Dr. Reinhard Fässler, PMID: 26821125) were cultured at 37 °C and 5% CO_2_ in Dulbecco’s modified Eagle’s medium (DMEM; Sigma-Aldrich, D6546), supplemented with 10% fetal bovine serum (FBS, Sigma-Aldrich, F2442), 4 mM L-Glutamate (Sigma-Aldrich, G7513), and 20 μg/mL gentamicin (Sigma-Aldrich, G1397). Cells were split every 2 days at a volume ratio of 1:5.

### Folding and purification of FluoroCubes

A modified Ref.^42^ protocol was employed to prepare the DNA FluoroCubes. In summary, the four 32 bp oligonucleotide strands (Integrated DNA Technologies, IDT) were dissolved in deionized water to a final concentration of 100 μM. Each strand was modified with biotin, Cy5, or both, with the sequence available in the supplementary data (Table S2). For a 50 μl folding reaction of the desired six-dye FluoroCubes, 5 μl of each of the four oligos were added to a PCR tube to reach the final concentration of 10 μM, filled with 25 μl of deionized water, and 5 μl of the 10x folding buffer (50 mM Tris pH 8.5, 10 mM EDTA and 400 mM MgCl_2_). The samples were placed in a thermocycler to fold the six dye FluoroCubes with the following temperature ramp: Denaturation occurred at 85 °C for 5 min, followed by cooling from 80 °C to 65 °C with a decrease of 1 °C per 5 min; Further cooling was occurred from 65 °C to 25 °C with a decrease of 1 °C per 20 min; The final holding temperature was at 4 °C. Folded products were analyzed by 2.0% agarose gel electrophoresis in TBE buffer (45 mM Tris-borate and 1 mM EDTA) with 12 mM MgCl_2_ at 70 V for 3 hours. To prevent potential unfolding, the gel box was placed in an ice bath during analysis. After completion of the gel run, it was analyzed using a custom-built system consisting of a 625 nm LED (ThorLabs, M625L3) and a bandpass filter (ET700/75, Chroma). DNA staining dyes such as ethidium bromide are prohibited from use as they will bind to the DNA and, therefore, might interfere with the absorbance and emission pattern of the FluoroCubes in downstream imaging applications. The desired band was then extracted from the gel using a razor blade. Afterward, the excised band was placed in a Freeze ‘N Squeeze™ DNA Gel Extraction Spin Column (BioRad, 732-6165) and centrifuged for 3 min at 12,000 rcf at 4 °C. The flow-through contained the desired FluoroCubes and was stored at 4 °C for a couple of weeks.

### PDMS gel substrate preparation for TFM

Employed glass coverslips (Marienfeld, 60 × 24 mm, #1.5H) underwent a thorough cleaning process in two steps. Initially, they were sonicated in a 5% v/v solution of Hellmanex III for 10 minutes, followed by sonication in 100% Ethanol. During each step, the coverslips were rigorously washed in double-distilled water for 5 minutes. Finally, compressed nitrogen was used to dry the coverslips.

Silicone substrates were fabricated following previous studies, combining a 1.2:1 or 1:1 weight ratio of two components, Cy-A and Cy-B (Dow DOWSIL™ CY 52-276 Kit), resulting in substrate stiffness of approximately 7 or 12 kPa, respectively^45,48,86^. The mechanical properties of the gels, such as rigidity, can be tuned by varying the ratio of base to crosslinking agent, as well as the curing temperature and duration. For example, a mixture prepared at a 9:10 base-to-crosslinker ratio and cured at 70 °C for 30 minutes resulted in a gel with an elastic modulus of approximately 12.7 kPa^15^. The gel solution was mixed thoroughly, centrifuged, and degassed before pipetting 500 µl of it onto a glass coverslip. The solution was spin-coated for 90 seconds using a home-made spinner, with acceleration and deceleration occurring every 30 seconds, to create a smooth, uniform surface. The gels were cured either at 80 °C for 2 hours or 70 °C for 30 min, after which they were stored at RT. The measured thickness of the gels was ∼10 µm, and the stiffness of the gel was confirmed by atomic force microscopy (AFM, Methods, see Figures S8 and S9). To functionalize the silicone surface, the gels were treated with a solution of 7% (vol/vol) 3-aminopropyl trimethoxysilane (APTES) in pure ethanol for 2 hours. The gels were washed with pure ethanol, a 1:1 volume ratio of ethanol and PBS, and finally with PBS alone, each for three cycles.

As a cell culture and imaging chamber, an 8-well bottomless sticky-slide (Ibidi, 80828) was mounted on top of each PDMS substrate. After washing the chambers three times with PBS (Sigma-Aldrich, D8537), the surfaces were incubated with 0.5 g/L BSA-Biotin (ThermoFisher, 29130) in PBS for 2 hours at RT. This incubation time can be reduced to 30 minutes if desired. Subsequently, the chambers were washed three times with PBS before incubation with 0.5 g/L Neutravidin (ThermoFisher, 31050) in PBS for 30 min at RT. The surfaces were washed three times with PBS and incubated with purified FluoroCube-DNA solution, diluted to 1 nM in PBS, for 10 min at RT. Afterward, the prepared samples were washed at least three times in PBS. A solution of 40 nm, red-orange fluorescent (565/580) FluoSpheres carboxylate-modified microspheres (Invitrogen, F8794) at a dilution of 1:1,000 in PBS was mixed with a final concentration of 100 µg/mL EDC and pipetted onto the top surface of the gel. The solution was incubated for 20 min at RT and washed three times with PBS. Next, the gels were coated with a solution of 10 μg/mL RGD peptides with 100 μg/mL of EDC in PBS and incubated for 2 hours at RT or overnight at 4 °C. After incubating, the chambers underwent multiple washes, initially with PBS and then with the cell culture medium, before being seeded with cells. The versatility of the chambers was further evaluated in the presence of extracellular matrix proteins, specifically fibronectin and collagen. After storage in the dark at 4°C, the chambers were tested at one- and two-week intervals, and fluorescence signals remained stable.

### Validation of the substrate stiffness by AFM measurements

Force spectroscopy measurements were performed on a JPK NanoWizard 3 atomic force microscope (JPK Instruments) using biosphere MLCT-SPH-5UM cantilevers (Bruker), with a spring constant of 0.393 Nm^−1^. For typical measurements, the set-point force was set to 1 nN, the acquisition speed to 1 µm.s^−1^, and the z length to 2 µm. The force spectroscopy measurements of the gels were conducted in PBS buffer containing 1% SDS to minimize the impact of the strong adhesive forces between the cantilever and the gels^86^. Data analysis was done using JPK data processing software (Version 6.1.142, JPK Instruments). The Young’s modulus was obtained by fitting the extended part of the force curves (at least 224) with a simple Hertz–Sneddon model, considering a spherical tip shape and a radius of 5 µm. The obtained Young’s modulus values were further processed in GraphPad Prism 10.4.0. This consisted of identifying and removing outliers and presenting the data as histograms for each elastomer variant (Figure S8).

### Microscopy

All microscopy data were acquired using an inverted IX83 microscope (Olympus, Japan) with an Abbelight SAFe 360 nanoscopy module, Oxxius lasers (LPX-640-500-CSB, Oxxius, France), and using the NEO Analysis software (Abbelight, France). The Abbelight setup enabled precise adjustment of the illumination angle, allowing smooth switching between EPI, HILO, and TIRF modes. Both TIRF and HILO illumination were tested and validated on PDMS substrates, yielding improved signal-to-noise ratios. To excite Cy5-FluoroCubes, 40 nm beads, and GFP-labeled proteins, the 640, 532, and 488 nm laser lines were integrated into the microscope through the ASTER technology with TIRF illumination mode. The illuminated lights were reflected onto the objective (100x oil-immersion objective, NA 1.5) with a quad-band dichroic mirror (Di03-R405/488/532/635-t1-25×36, Semrock, USA). The emitted light was initially filtered through a quadband bandpass filter (FF01-446/510/581/703-25 BrightLine quad-band bandpass filter, Semrock, USA) and subsequently with a single bandpass filter (F-445/45, F-510/20, F-582/64 Semrock, USA), which were mounted on an Optospin fast filter wheel (Cairn, England). The fluorophore emissions were recorded with a 2048 × 2048-pixel sCMOS camera (Orca-fusion C1440-20UP, Hamamatsu, Japan) with an optical pixel size of 97 nm. Before imaging, the cells were washed twice with DMEM media and then switched to the imaging buffer, which includes DMEM media with the final concentration of 0.56 mg/mL of glucose oxidase (Aspergillus niger-Type VII, lyophilized powder, Sigma-Aldrich, G2133) and 34 μg/mL of catalase (bovine liver -lyophilized powder, Sigma-Aldrich, C40) alongside Trolox ((+/−)-6-hydroxy-2,5,7,8-tetra-methylchromane-2-carboxylic acid, Sigma-Aldrich, 238813-5G). The 100× Trolox stock solution was prepared by combining 100 mg of Trolox, 430 μl of methanol, and 345 μl of 1 M NaOH in 3.2 ml of H₂O. Image acquisition of FluoroCubes and fluorescent beads were performed over 100 frames and 50 frames, respectively. Fluorescent imaging was performed in TIRF mode using 20% laser power for the 532 nm and 640 nm lines (maximum laser power of 90 and 228 mW at the optical fiber output, respectively), with exposure times of 50 ms. Images were acquired over a 928 × 928 pixel field of view, corresponding to approximately 90 × 90 μm. After the cells were detached using 0.5% Trypsin-EDTA (Gibco, 15400054), the imaging media was exchanged to capture the gel’s relaxed state. Confocal images of the supplementary data were conducted on a STED microscope (Abberior Instruments GmbH, Göttingen) equipped with a 100x oil-immersion objective, NA 1.4. Imaging parameters included voxel dimensions of 100 × 100 × 200 nm, dwell time of 5.000 μs, and a line accumulation factor of 4 to enhance signal quality. Subsequent image processing and analysis were carried out using Fiji and MATLAB, ensuring consistent and reproducible data presentation.

### Beads and FluoroCubes surface stability and homogeneity

To compare the surface stability and cellular uptake of beads versus FluoroCubes, cells were seeded on substrates functionalized with beads and/or FluoroCubes. After 16 hours of incubation, cells were washed three times with phosphate-buffered saline (PBS) to remove unbound particles and then fixed with 3.7% formaldehyde (Carl Roth, 10.1) in PBS for 20 minutes at RT. Following fixation, samples were washed three times with PBS and imaged in our imaging buffer using confocal microscopy. To assess surface homogeneity for beads and FluoroCubes in the presence of cells, we computed a normalized coefficient of variation (CV) that quantifies local intensity variation under the cell relative to outside the cell. Each cell was segmented using a polygonal ROI. The image was then divided into non-overlapping 32×32-pixel tiles, and the mean intensity of each tile was calculated (excluding pixels that are masked by the ROI). For a region R, the CV was defined as 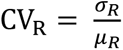, where *σ*_*R*_ is the standard deviation of the tile-mean intensities and *μ*_*R*_ is their average. The normalized CV was computed as the ratio 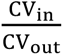, where “in” refers to the region beneath the cell footprint and “out” refers to the extracellular region outside the cell. Higher values indicate greater local intensity variation under the cell relative to outside, i.e., a less uniform probe distribution.

### Photostability

FluoroCubes and 40 nm dark red (660/680) FluoSphere beads were excited in the same channel and imaged under identical conditions in optimized imaging buffer. Imaging was carried out under TIRF illumination for 10,000 consecutive frames, using a 200 ms exposure time and 20% laser power. Localization and photon number estimation were performed using the Abbelight NEO software suite. Photon counts from each localized event were summed per frame and normalized to the maximum photon count in the time series. All subsequent computation and fitting analyses were carried out in MATLAB.

### Quantifying substrate displacements with optical flow tracking

Displacement fields were acquired by comparing images of different fiducial markers before and after detachment of the cells by trypsinization. We employed the standard Kanade-Lucas-Tomasi (KLT) optical flow feature tracking algorithm^35,38^ to track the displacements locally. The standard KLT algorithm effectively minimizes the sum of squared intensity differences in an interrogation window between consecutive frames with a local displacement vector ***u*** = (*u*_*x*_, *u*_*y*_),

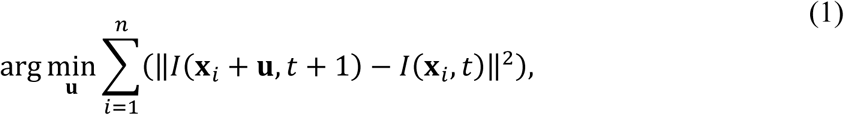

where *I*(**x**_*i*_, *t*) denotes the pixel intensity at position **x**_*i*_ and time *t*. Linearizing *I*(**x**_*i*_ + **u**, *t* + 1) for small displacements yields the standard optical-flow equation in the form of ***Au*** = −***b***, where ***A*** and ***b*** are functions of spatial and temporal intensity gradients (Equation S2). To extend the KLT algorithm for dual channels, we modified the expression of ***A*** for a second channel using a cross-correlation-based weighting scheme (Equation S8-10), which effectively imposes a stronger constraint on the optimization problem described by Equation 1. By this modification, we were able to track the coherent motion of the two channels simultaneously.

### Synthetic TFM images

The performance of the image analysis procedures was assessed with synthetic TFM data. We created 20 elliptical traction patches that are evenly distributed around the periphery of a circular cell with a radius of 8 *μm*. Each traction patch has a major axis with length 1 *μm* and a minor axis with length of 0.25 *μm*, where the major axis points toward the cell center. Traction forces inside each patch point toward the cell center and have a constant magnitude of 500 Pa. Assuming a Young’s modulus of 3000.00 Pa and a Poisson’s ratio of 0.5, we calculated the exact displacement vectors on a grid with 20 nm spacing. Next, we randomly selected a number of grid points as reference positions for fluorescent labels, ensuring a label density of approximately 5 *μm*^−2^. A Gaussian point spread function with a standard deviation of 2 pixels was applied to each reference position to generate images. A deformed bead image was acquired similarly, with the new label positions computed based on the previously obtained displacements.

We generated two sets of reference-deformed image pairs for each of the two imaging channels. Note that neither background noise nor local variations in the PSF are included in the synthetic data for the *in silico* assessment of the modified optical flow tracking algorithm’s improvement.

Consequently, we only assessed tracking errors resulting from the high density of beads with overlapping point spread functions.

### Image preprocessing

Image processing was performed using MATLAB. The employed functions are written in *italic* below. All raw images of fiducial markers were preprocessed to enhance features, using a difference-of-Gaussian filter with a filter size of 5 pixels in both dimensions. The standard deviations of the Gaussian filters varied depending on the image qualities and were set to 1.5 and 2.5 pixels, respectively. The *imgaussfilt* function was employed for filtering, and the *imadjust* was applied for contrast adjustment. Subsequently, marker detection and tracking were done with feature-enhanced images. Before analysis, we corrected for a stage shift using *normxcorr2* with subpixel accuracy achieved by fitting the correlation peak with a 2D Gaussian function, implemented with *lsqcurvfit*. After aligning the images, we used *vision.LocalMaximaFinder* or *detectMinEigenFeatures* to determine pixel positions of feature points. For conventional KLT tracking, the positions of the detected fiducials were then fed into *vision.PointTracker*.

### Assessment of image quality and point spread function using fluorescent beads

A customized MATLAB script was used to calculate the signal-to-background ratio and fluorescence intensity profile around every nanobead, as shown in Figure 1. First, the beads were localized using the function *vision.LocalMaximaFinder*. Subsequently, an interrogation window of 7-by-7 pixels was placed on top of every local fluorescence intensity maximum to extract images of the beads. Additionally, images of the background signal were acquired by excluding the extracted bead images through masking. Then, the signal-to-background ratio was computed by averaging the local maximal intensities divided by the average intensity of the background pixels. The average fluorescence intensity profile was computed by averaging the intensities along the horizontal center line for all extracted bead images. For better visualization, the profile was fitted with a Gaussian function. The intensities were normalized by a constant.

### Traction field reconstruction

Once the displacement field was obtained, we used BFTTC to reconstruct the traction field with an optimal regularization parameter^41^. This parameter was estimated based on the noise level in the displacement field. In our implementation, displacement vectors from regions near the boundaries were always considered, as these areas are far from cells and are not expected to deform. To account for the finite thickness of our substrates (∼10 µm), we employed a modified Green’s function^69–72^. The algorithm assumes a purely elastic substrate; Although PDMS exhibits some viscous creep, the thin, stiff CY52-276 gels used here minimize time-dependent deformation, making viscous effects negligible on the timescale of our experiments^87,52^.

### Displacement correlation between two channels

To quantify the consistency of the displacement fields from both probes, we used bilinear interpolation to project the measured displacement vectors onto a common regular grid for the two individual channels. We then mapped the interpolated vector components from the bead channel to the same components from the FluoroCube channel based on their positions on the grid. The correlation between the two channels was visualized by plotting the FluoroCube displacement as a function of the corresponding bead displacements. A slope of 1 would be expected for perfect correlation.

## Supporting information

Comparison of bead and FluoroCube imaging results on PDMS across different illumination modes; Quantification of the deviation of traction magnitude f

## Code availability

The dual-channel KLT tracking code and representative example datasets used in this study are hosted on GitHub at https://github.com/CellMicroMechanics/Dual_channel_KLT_tracking_for_TFM

## Supporting Information

Comparison of bead and FluoroCube imaging results on PDMS across different illumination modes; Quantification of the deviation of traction magnitude for simulated data; Effect of noisy images on traction reconstruction with optical-flow tracking; Reduction of fluctuation-induced noise by averaging successive frames; Images of fluorescent probes from individual channels acquired by using different approaches; Validation of tracking algorithms with controlled stage-shift experiments; Comparison of measured displacement fields from the two individual channels and the combined two channels using a very small, 10-pixel-sized tracking window; AFM measurement of the mechanical stiffness of the TFM substrates; Distribution of the thickness of PDMS gels; Demonstration of TFM with FluoroCubes and fluorescent beads as fiducial markers; Parameter values used for the generation of data shown in Figure 3 and 4 of the main text; Oligonucleotides used in this study and their respective sequences and modifications; Tested STORM buffers; Cy5 FluoroCubes in STORM Buffer; Cy5 FluoroCubes in Optimized Buffer; 40 nm Red-Orange (565/580) FluoSpheres in STORM Buffer; Derivation of the modified Kanade-Lucas-Tomasi (KLT) optical flow tracking algorithm.

## Author contributions

AM, PL, and SG performed experiments; JJ wrote the code. AM, JJ, SG, and BS analyzed the data; DAH, RS, and MS provided fluorescent constructs. AM and JJ contributed equally. All authors participated in writing the manuscript.

## Acknowledgments

The authors gratefully acknowledge the generous gift of Kind^Ko+K2GFP^ cells from Prof. Dr. Reinhard Fässler (MPI of Biochemistry, Munich, Germany) and the gift of constructs and cell lines from Prof. Dr. Carsten Grashoff (University of Münster, Germany). They also thank Prof. Dr. Markus Meissner (Experimental parasitology, LMU, Munich, Germany) for generously providing access to the confocal microscopy and Dr. Javier Periz for his invaluable support and guidance with the device. AM, JJ, and BS received funding from the European Research Council (ERC) under the European Union’s Horizon 2020 research and innovation program (BacForce, G.A.No. 852585). JJ and BS acknowledge funding by the Deutsche Forschungsgemeinschaft (DFG, German Research Foundation), Project no 492014049. DAH, RS, and MS received funding from ERC under the European Union’s Horizon 2020 research and innovation funding program (G.A.No. 835102).

