## Supplementary material for "Traction Force Microscopy with DNA FluoroCubes": Comparison of bead and FluoroCube imaging results on PDMS across different illumination modes; Quantification of the deviation of traction magnitude f

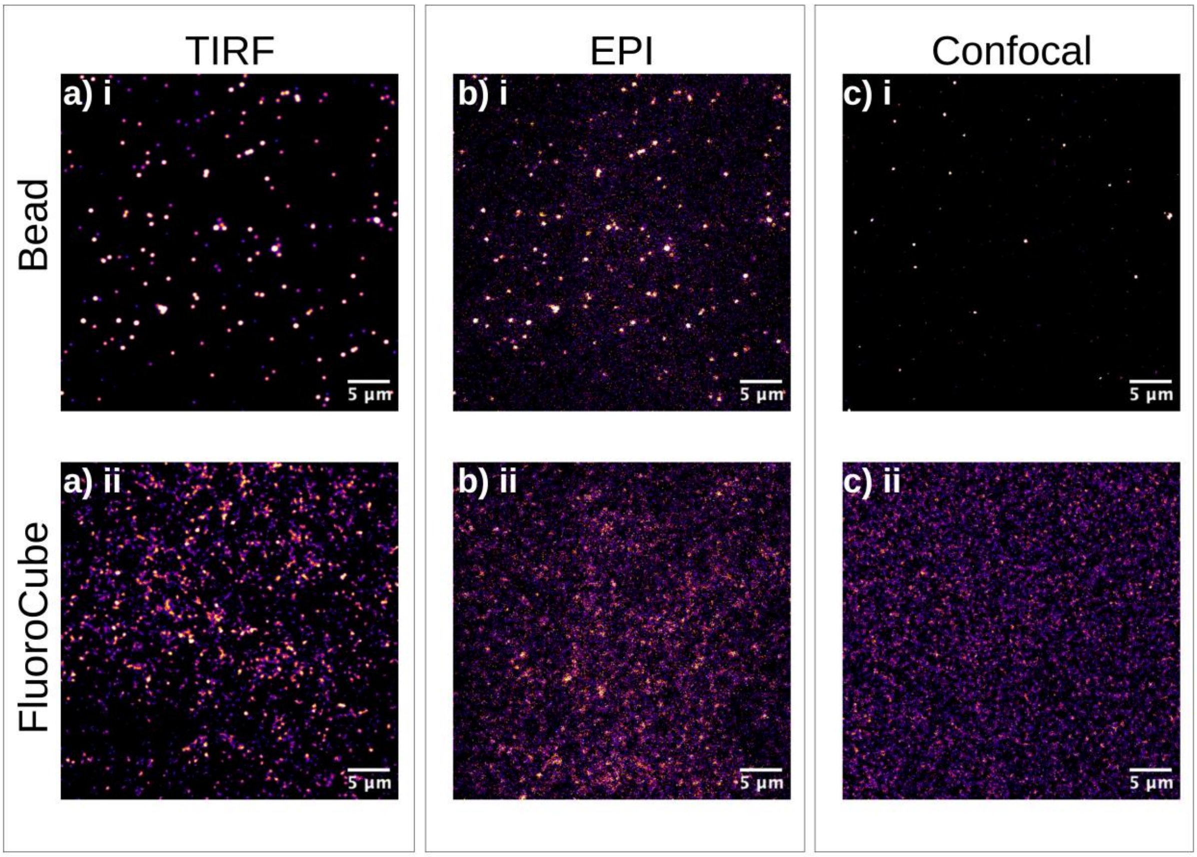


**Figure S1.** Comparison of bead and FluoroCube imaging results on PDMS across different illumination modes. (a-c) i In the bead channel, confocal microscopy exhibits an excellent signal clarity. (a-c) ii In the FluoroCube channel, differentiation between signal and background noise is challenging in both epifluorescence and confocal modes. However, TIRF microscopy allows one to clearly resolve FluoroCubes, offering significantly enhanced signal discrimination compared to the two other imaging techniques.

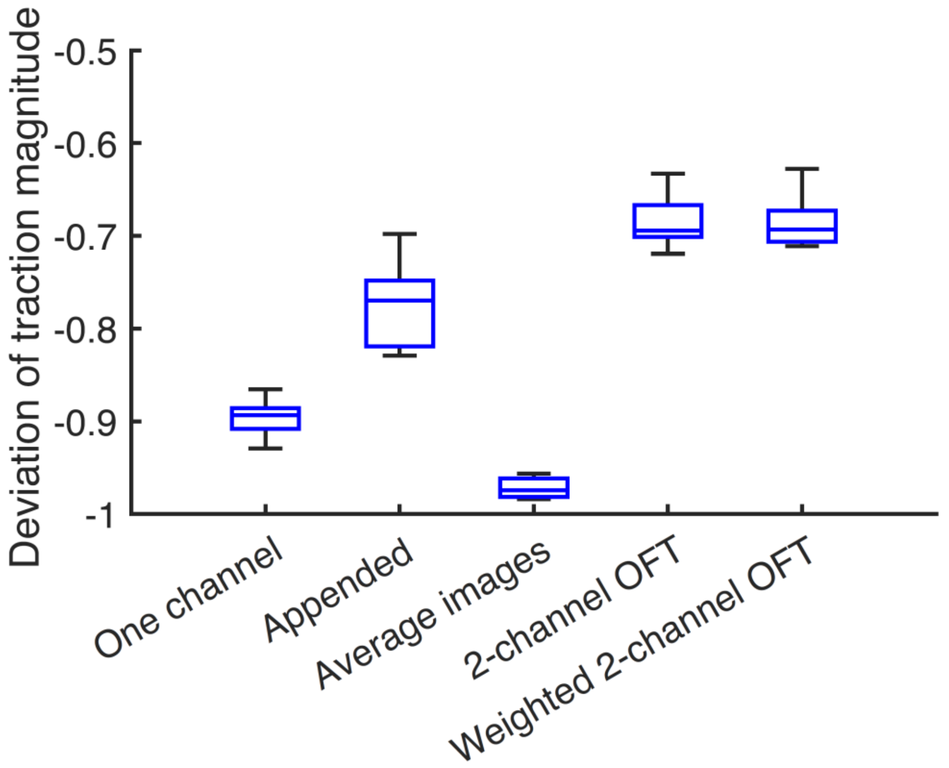


**Figure S2.** Quantification of the deviation of traction magnitude for simulated data. The simulated dataset is the same as that used for Figure 3 in the main text. The quantification of deviation of traction magnitude is done for different tracking routines, i.e., traction reconstruction from one-channel conventional KLT (One channel), appending the KLT tracking results from the two channels (Appended), KLT tracking result from the averaged image of the two channels (Average images), the modified KLT tracking for the two channels (2-channel OFT), and the modified KLT tracking with cross-correlation-based weighting for the two channels (Weighted 2-channel OFT). This shows that, although systematic traction underestimation is likely always present, the combination of two image channels with optical flow tracking (OFT) yields the best results due to the high spatial resolution of this approach.

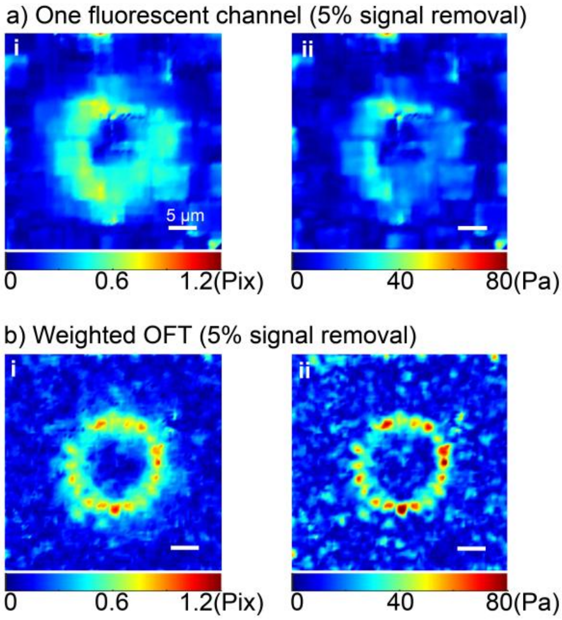


**Figure S3.** Effect of noisy images on traction reconstruction with optical-flow tracking. Synthetic traction force microscopy dataset incorporating intensities randomized uniformly in the range of 700~1500 photon counts and a 5% loss of fiducial markers to mimic photobleaching. The ground-truth traction pattern matches that shown in Figure 3a of the main text. (a) Displacement field (i) and reconstructed traction map (ii) obtained using the conventional single-channel KLT algorithm. (b) Displacement field (i) and corresponding traction map (ii) obtained using the modified dual-channel KLT algorithm.

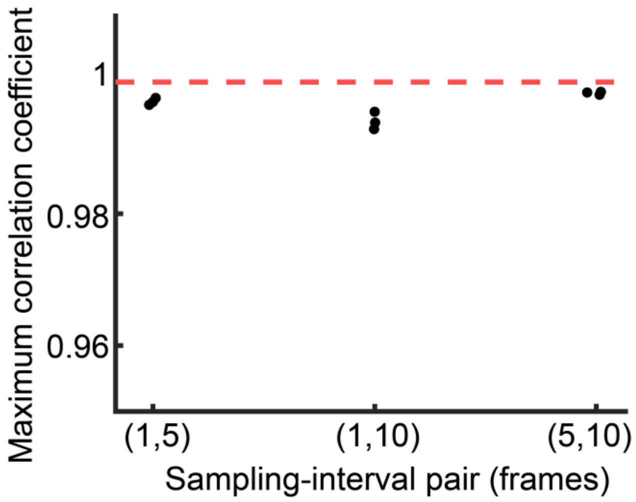


**Figure S4.** Reduction of fluctuation-induced noise by averaging successive frames. Maximum cross-correlation coefficients between images generated from different time-averaging intervals. Values near 1 indicate that averaged images are nearly identical across intervals. The high correlations demonstrate that time averaging effectively suppresses intensity-fluctuation noise, providing stable inputs for displacement tracking.

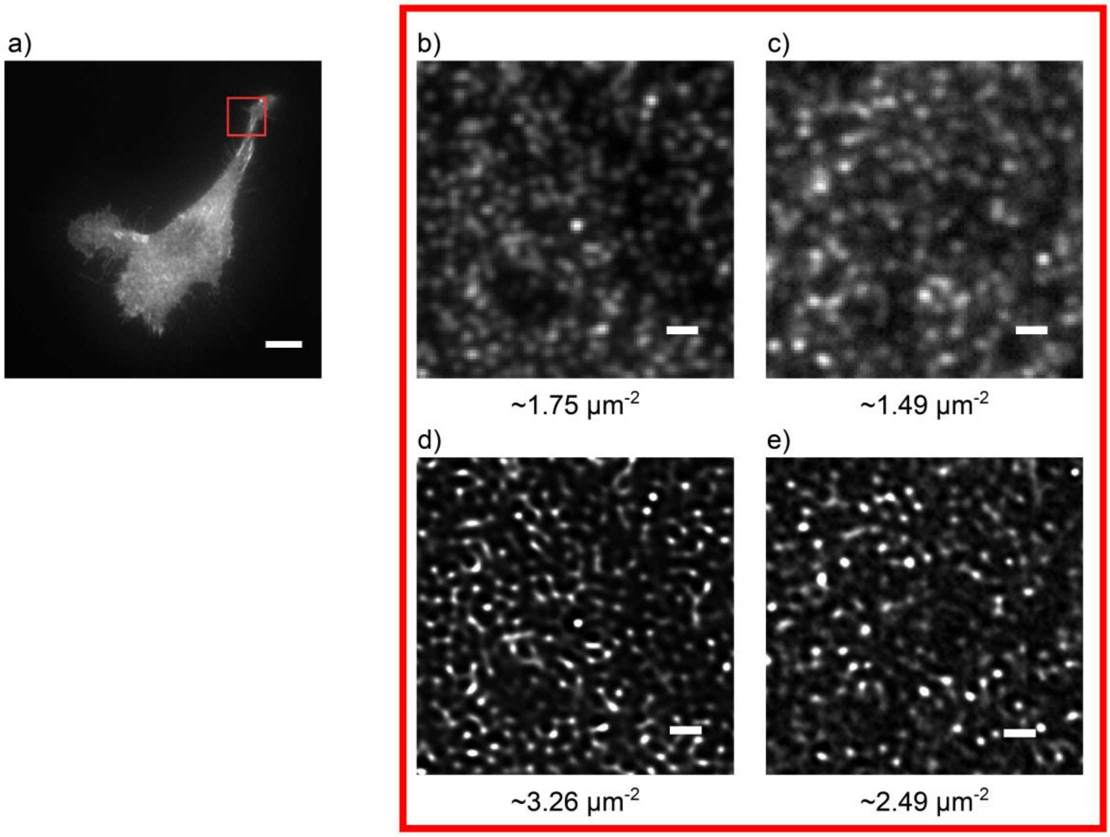


**Figure S5.** Images of fluorescent probes from individual channels acquired by using different approaches. (a) A murine kidney fibroblast cell with GFP-labeled kindlin-2 (KindKo+K2GFP cell line), scale bar is 10 μm. (b) and (c) Widefield images of 40 nm fluorescent beads and FluoroCubes acquired in TIRF mode, respectively. (d) and (e) Reconstructed images of beads and FluoroCubes, respectively, via super-resolution imaging based on autocorrelation with two-step deconvolution (SACD)[1]. Scale bars are 1 μm. The numbers indicate the estimated probe number density.

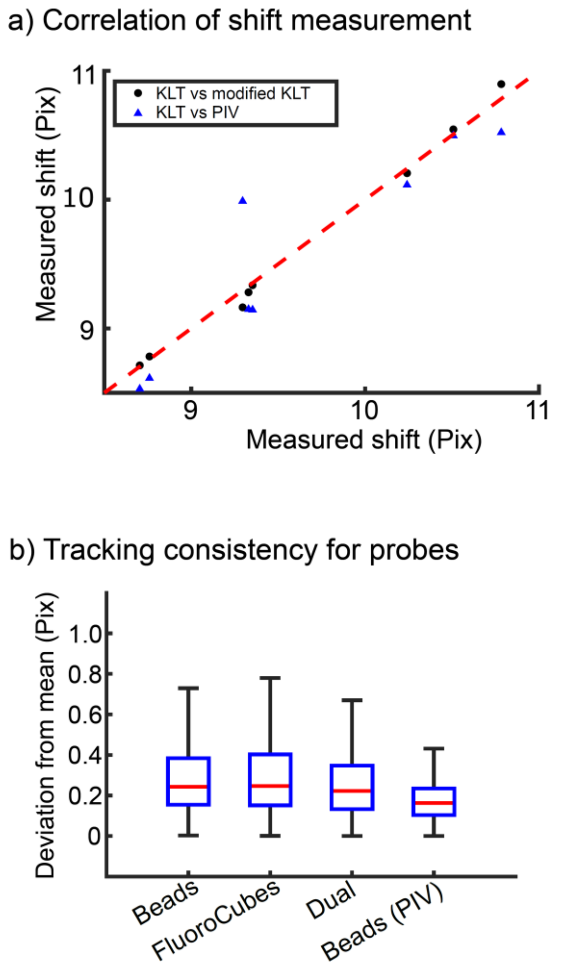


**Figure S6.** Validation of tracking algorithms with controlled stage-shift experiments. (a) Correlation between displacement fields obtained with the standard KLT algorithm and those from either the modified KLT or standard PIV methods. (b) Mean deviation of the tracked shifts at individual detected points for each method. Smaller deviations indicate greater tracking stability in recovering uniform stage movements.

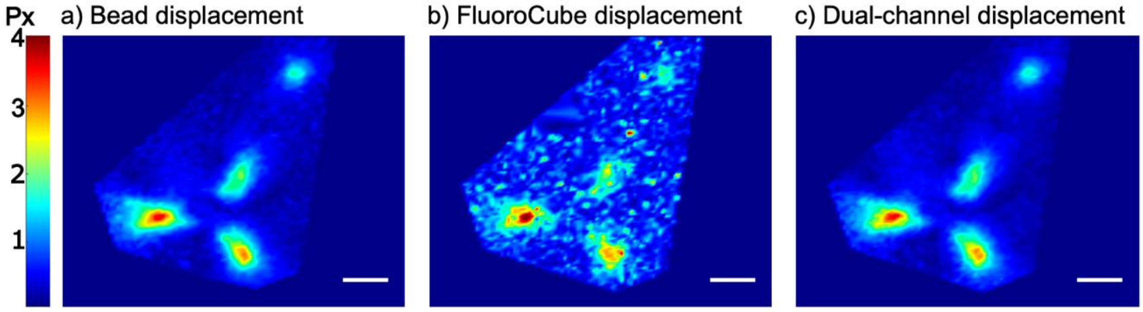


**Figure S7.** Comparison of measured displacement fields from the two individual channels and the combined two channels using a very small, 10-pixel-sized tracking window. For such a small window size, the tracking error in the denser FluoroCube channel is significant because the information contained in the small window is frequently insufficient to determine its precise location in the displaced frame of reference. When the two channels are combined, the additional information contained in the bead channel improves the result, and additional details of the displacement field that are potentially only visible in the FluoroCube channels are integrated. All scale bars are 12.5 μm.

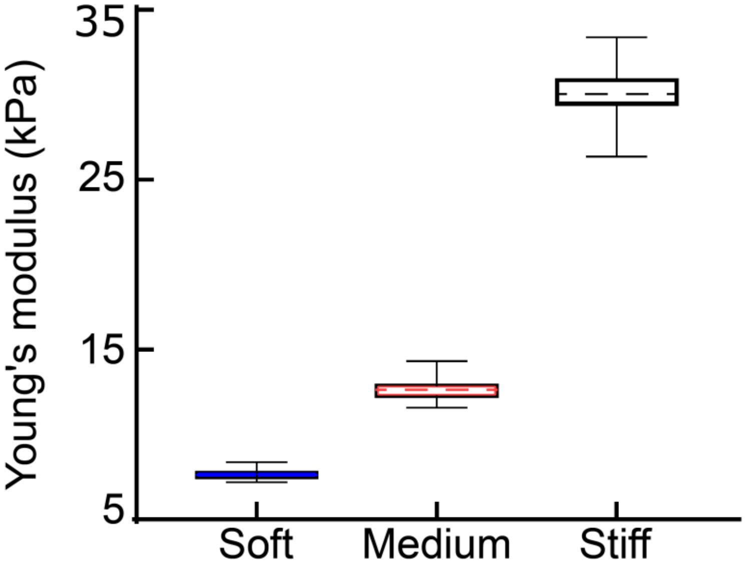


**Figure S8.** AFM measurement of the mechanical stiffness of the TFM substrates. Box plots of Young's moduli were derived from AFM measurements for three ratios of Dow DOWSIL™ CY 52-276 PDMS gels, prepared with varying base elastomer-to-curing agent ratios. The mean values for the three different samples are 7.624 (±0.197) kPa, 12.589 (±0.448) kPa, and 30.141 (±1.014) kPa, respectively.

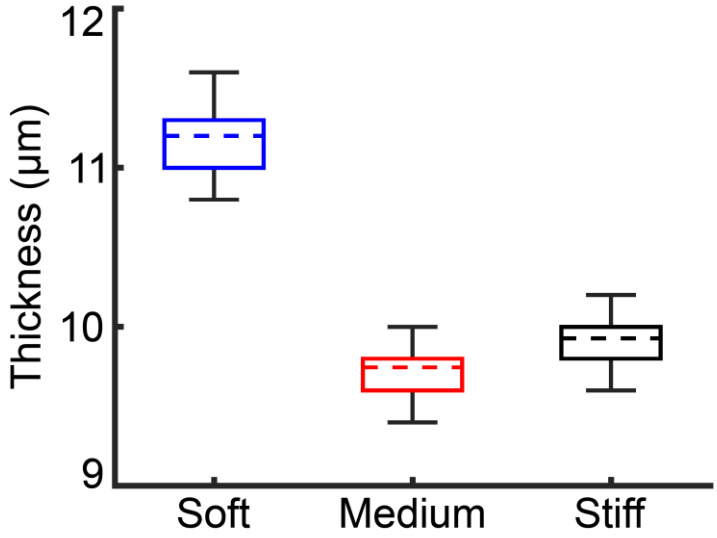


**Figure S9.** Distribution of the thickness of PDMS gels. The thickness was measured using confocal microscopy as follows. A 1:40,000 dilution of 100 nm red-orange fluorescent carboxylate-modified microspheres in ethanol was spin-coated onto a cover glass, followed by coating with the gel pre-polymer. After polymerization, the gel surface was functionalized with APTES to facilitate the conjugation of fluorescent microspheres (1:20,000 in PBS). Z-stack images were acquired at 200 nm intervals across 20 distinct regions of the gel. Thickness was determined by identifying the z-positions of the two highest fluorescence intensity peaks in the histogram, corresponding to the cover glass surface and the gel-medium interface, and calculating the difference between these values. The stiffness corresponds to the measurement shown in Figure S5. The mean thickness in each sample is listed as follows: 11.17 (±0.20) µm for the soft sample, 9.75 (±0.43) µm for the median stiff sample, and 9.93 (±0.15) µm for the stiff sample.

**Figures S10-12** demonstrate the robustness of our approach by showing results for different cells.

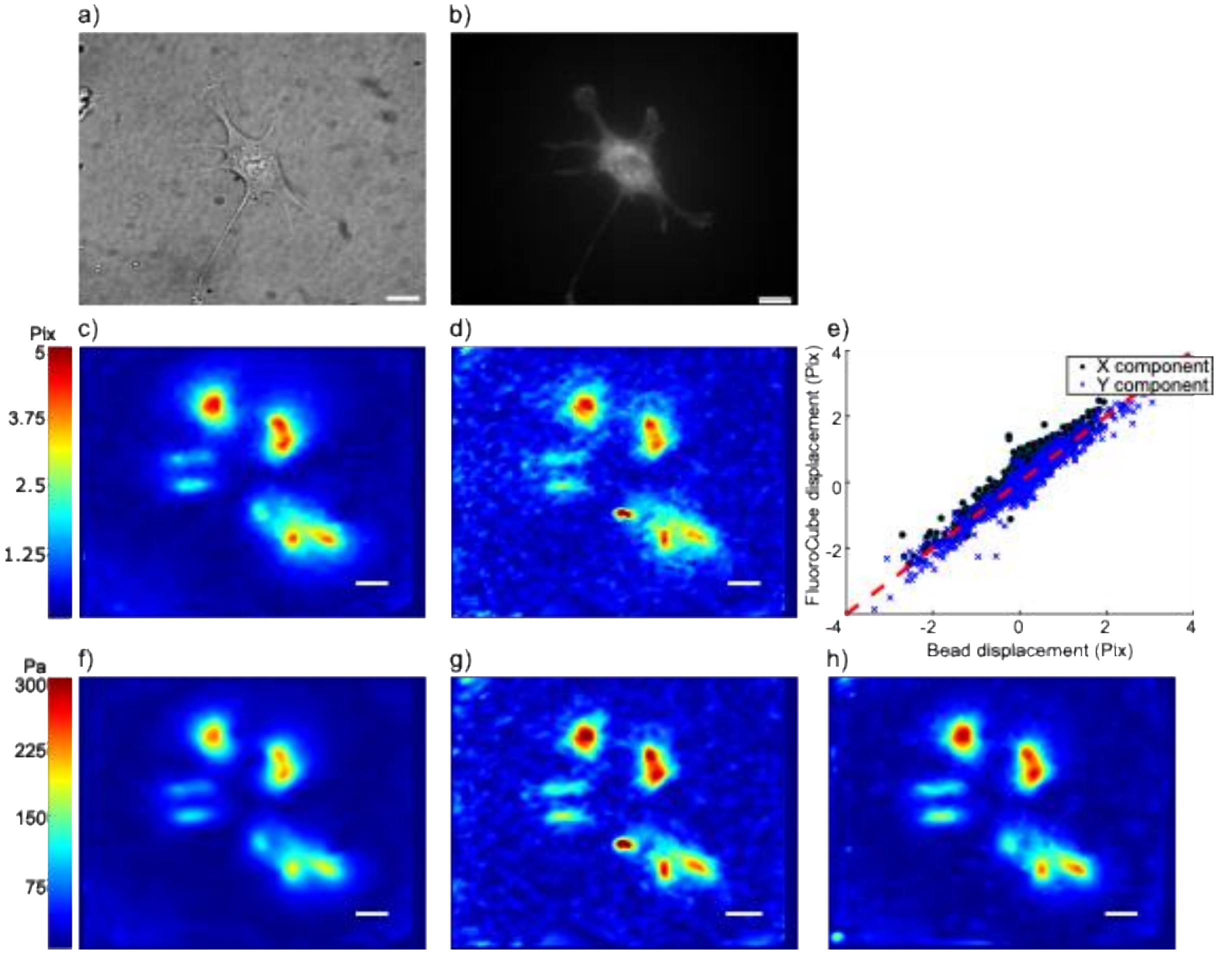


**Figure S10.** Demonstration of TFM with FluoroCubes and fluorescent beads as fiducial markers. (a) and (b) Bright field image of a murine kidney fibroblast with GFP-labeled kindlin-2 (Kind$^{Ko+K2GFP}$ cell line) and the corresponding fluorescence image; (c) and (d) Displacement fields measured in the bead and FluoroCube channel, respectively; (e) Displacement correlation between the two channels in the experimental images; (f) Traction forces calculated from the displacements in (c) with BFTTC; (g) Traction forces calculated from the displacement in (d); (h) Traction forces calculated from dual-channel optical flow tracking results. All scale bars are 12.5 μm. The Young's modulus of the gel stiffness is 12 kPa.

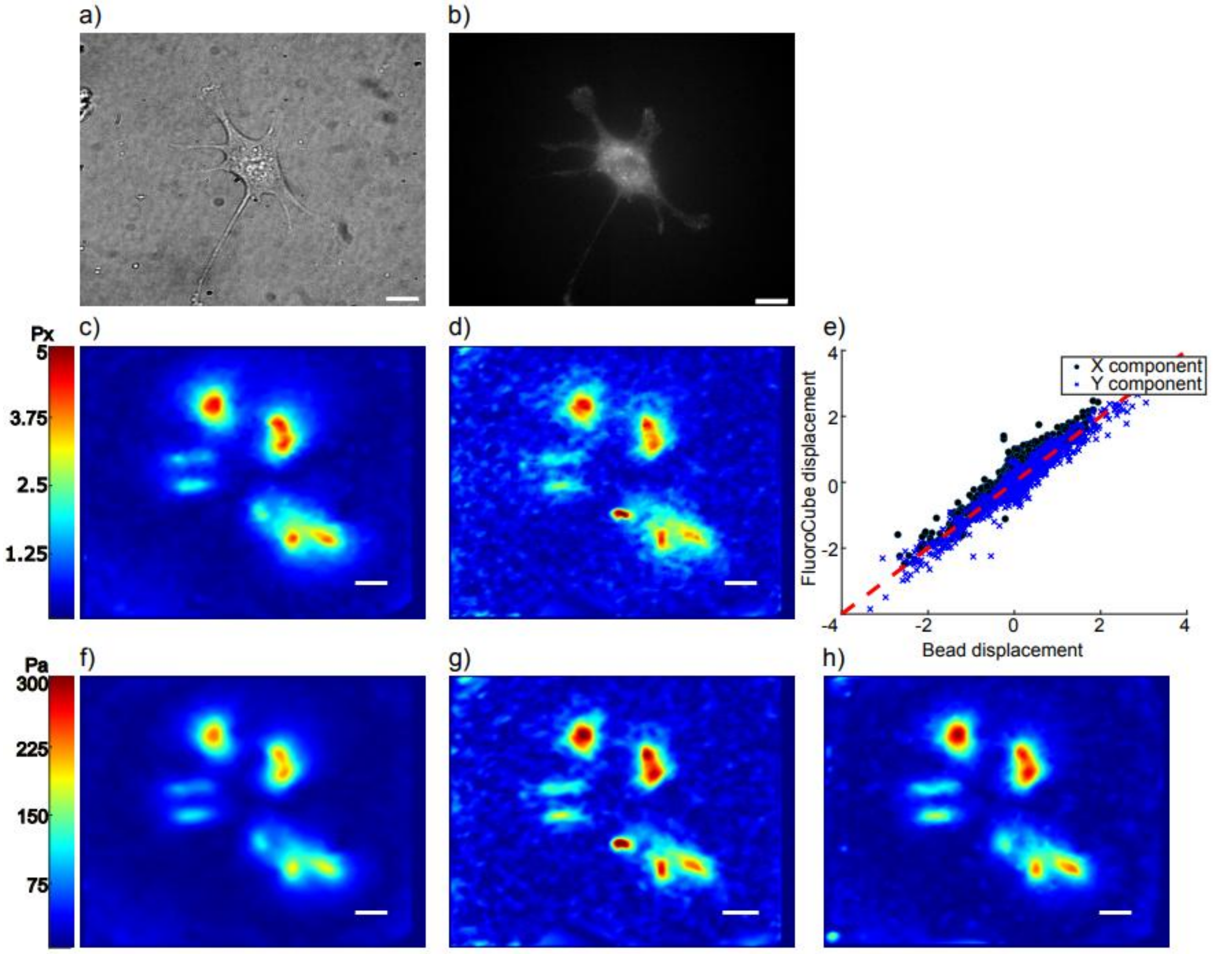


**Figure S11.** Demonstration of TFM with FluoroCubes and fluorescent beads as fiducial markers. (a) and (b) Bright field image of a murine kidney fibroblast with Ecto-tag β1 integrins expressing GFP and the corresponding fluorescence image; (c) and (d) Displacement fields measured in the bead and FluoroCube channel, respectively; (d) Displacement correlation between the two channels; (f) Traction forces calculated from the displacements in (c) with BFTTC; (g) Traction forces calculated from the displacement in (d); (h) Traction forces calculated from dual-channel optical flow tracking results. All scale bars are 12.5 µm. The Young's modulus of the gel stiffness is 12 kPa.

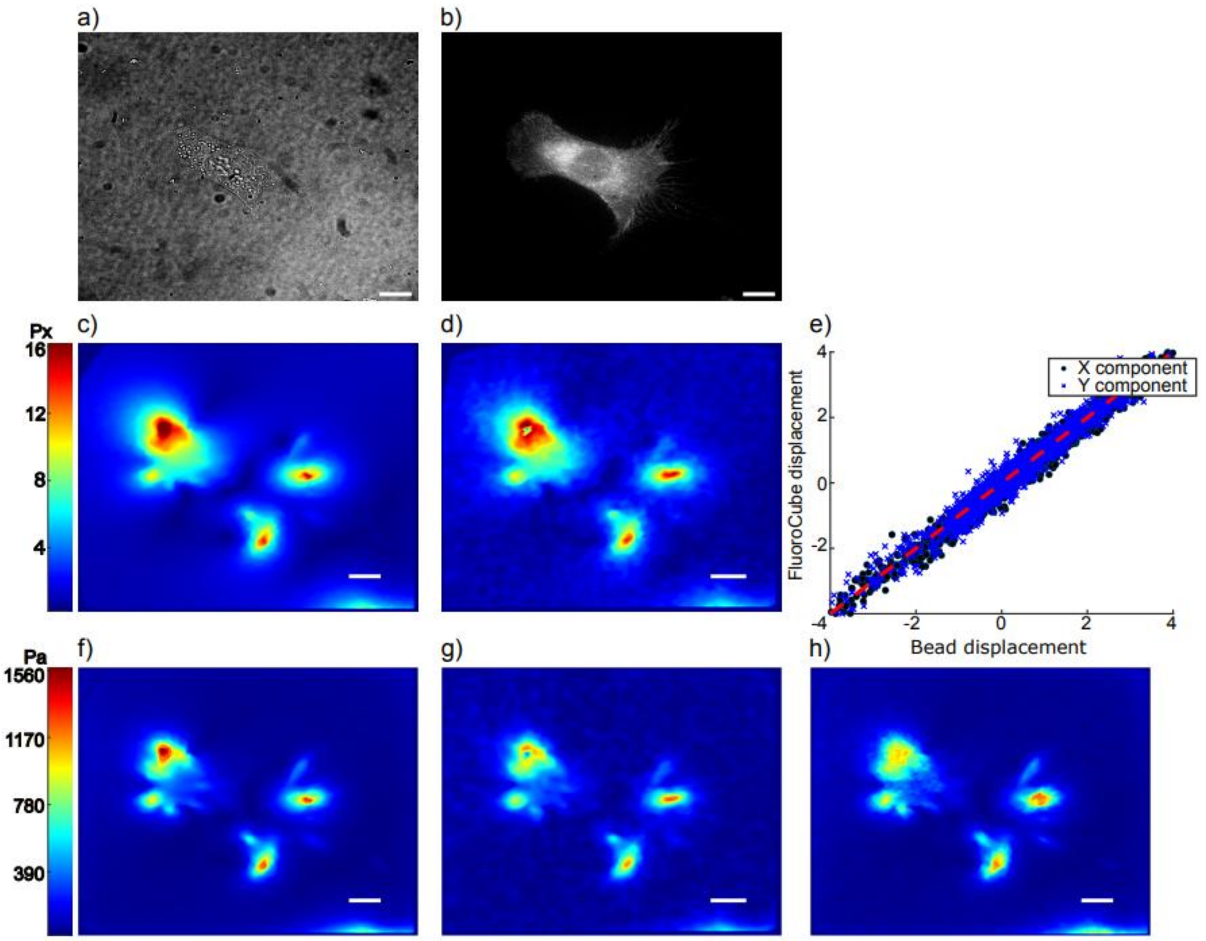


**Figure S12.** Demonstration of TFM with FluoroCubes and fluorescent beads as fiducial markers. (a) and (b) Bright field image of a murine kidney fibroblast with Ecto-tag β1 integrins expressing GFP and the corresponding fluorescence image; (c) and (d) Displacement fields measured in the bead and FluoroCube channel, respectively; (d) Displacement correlation between the two channels; (f) Traction forces calculated from the displacements in (c) with BFTTC; (g) Traction forces calculated from the displacement in (d); (h) Traction forces calculated from dual-channel optical flow tracking results. All scale bars are 12.5 μm. The Young's modulus of the gel stiffness is 12 kPa.

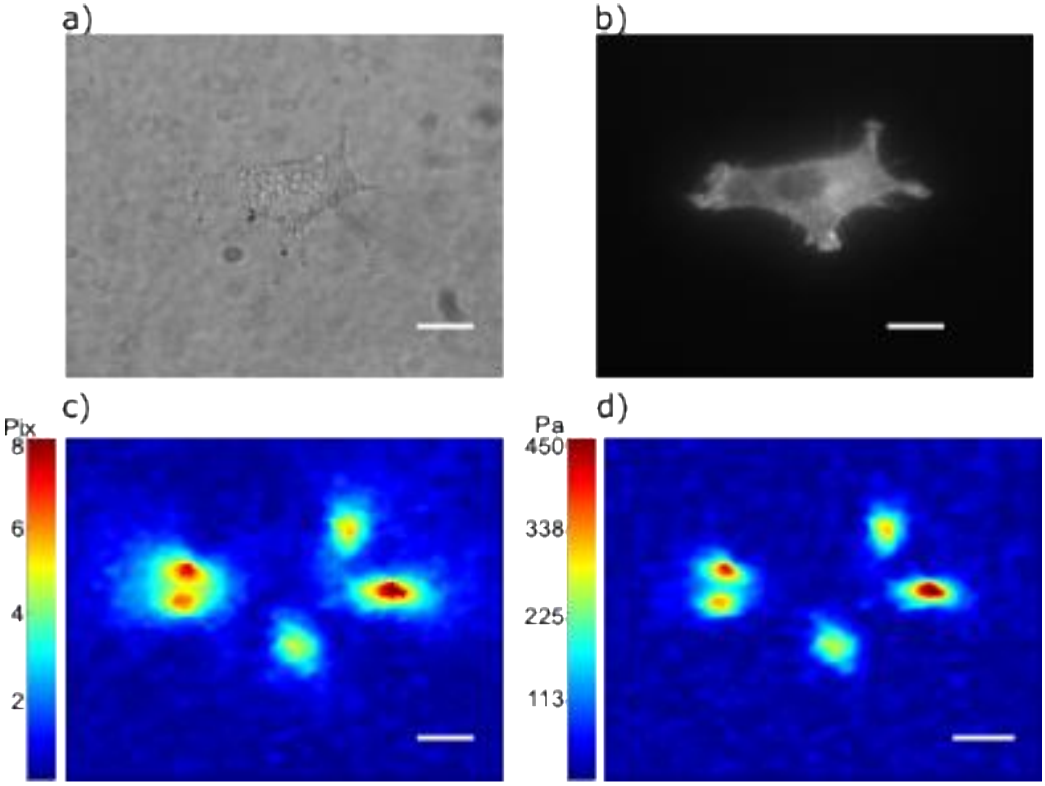


**Figure S13**. Demonstration of TFM with only FluoroCubes as fiducial markers. (a) and (b) Bright field image of a murine kidney fibroblast with Ecto-tag β1 integrins expressing GFP and the corresponding fluorescent image; (c) Displacement field measured from the FluoroCube channel; (d) Traction forces calculated from the displacement in (c) with BFTTC. All scale bars are 12.5 □m. The Young's modulus of the gel is 7 kPa.

**Table S1.** Parameter values used for the generation of data shown in Figures 3 and 4 of the main text. The analysis involved the tracking of fiducials and traction reconstruction with Bayesian Fourier Transform Traction Cytometry (BFTTC).

<table>
<tr><td colspan="6">Figure 3 parameters</td></tr>
<tr><td colspan="2">Vision.PointTracker (Matlab built-in)</td><td colspan="2">Modified KLT tracking</td><td colspan="2">FTTC parameters</td></tr>
<tr><td>Image pyramid levels</td><td>3</td><td>Image pyramid levels</td><td>3</td><td>Young’s modulus (Pa)</td><td>3000</td></tr>
<tr><td rowspan="2">Maximal number of iterations</td><td rowspan="2">30</td><td rowspan="2">Maximal number of iterations</td><td rowspan="2">30</td><td>Poisson’s ratio</td><td>0.5</td></tr>
<tr><td>Micron per pixel</td><td>0.1</td></tr>
<tr><td rowspan="2">Interrogation window size (pixel)</td><td rowspan="2">17-by-17</td><td rowspan="2">Interrogation window size (pixel)</td><td rowspan="2">17-by-17</td><td rowspan="5">Optimal normalized regularization parameters (pix$^2$) computed by BFTTC software.</td><td>815.3439 (Fig 3b)</td></tr>
<tr><td>728.3066 (Fig 3c)</td></tr>
<tr><td rowspan="3">Maximal bidirectional error (pixel)</td><td rowspan="3">1.5</td><td rowspan="3">Convergence (pixel)</td><td rowspan="3">0.01</td><td>214.0196 (Fig 3d)</td></tr>
<tr><td>33.1114 (Fig 3e)</td></tr>
<tr><td>22.3688 (Fig 3f)</td></tr>
</table>

<table>
<tr><td colspan="6">Figure 5 parameters</td></tr>
<tr><td colspan="2">Minimum eigen feature detection</td><td colspan="2">Modified KLT tracking</td><td colspan="2">FTTC parameters</td></tr>
<tr><td>Minimal quality</td><td>FluoroCubes: 0.0015<br>Beads: 0.0025</td><td>Image pyramid levels</td><td>2</td><td>Young’s modulus (Pa)</td><td>$10^4$</td></tr>
<tr><td>Gaussian filter size (pix)</td><td>5</td><td>Maximal number of iterations</td><td>100</td><td>Poisson’s ratio</td><td>0.5</td></tr>
<tr><td colspan="2" rowspan="5"></td><td rowspan="3">Interrogation window size (pixel)</td><td rowspan="2">31-by-31 Fig 4c) i,ii</td><td>Micron per pixel</td><td>0.1</td></tr>
<tr><td rowspan="2">Optimal normalized regularization parameters ($pix^2$) computed by BFTTC software.</td><td rowspan="2">FluoroCubes: 324.529<br>Beads: 86.927<br>Dual channel: 66.71</td></tr>
<tr><td>10-by-10 Figure S7c</td></tr>
<tr><td>Convergence (pixel)</td><td>0.01</td><td colspan="2" rowspan="2"></td></tr>
<tr><td>Number of weighting iterations</td><td>2</td></tr>
</table>

**Table S2.** Oligonucleotides used in this study and their respective sequences and modifications

| Name | Sequence (5´-3´) | Modification | | Vendor |
|---|---|---|---|---|
| | | 5´ | 3´ | |
| FC_SC_01 | ATGAGGTGTATGTGTAGAGTGATGGATGTAGT | Cy5 | Biotin | IDT |
| FC_SC_02 | AGGATGAGTGAGAGTGAGATGAGAGTAGATGT | Cy5 | Cy5 | IDT |
| FC_St_02 | CACTCTCACACCTCATACATCTACCATCACTC | Cy5 | Cy5 | IDT |
| FC_St_01 | TACACATACTCATCCTACTACATCTCTCATCT | Biotin | Cy5 | IDT |

**Tested STORM buffers.** Different dSTORM buffers, including DMEM (Sigma-Aldrich, D6546), Leibovitz's L-15 Medium without phenol red (Gibco,21083-027), DMEM without phenol red (Gibco, 21063-029), Abbelight's SMART kit buffer, Tris-STORM (50 mM Tris pH 8.0, 10 mM NaCl, 10% Glucose), and PBS-STORM (PBS pH 7.4, 10% Glucose). All buffers included the final concentration of 0.56 mg/mL of glucose oxidase (Aspergillus niger-Type VII, lyophilized powder, Sigma-Aldrich, G2133) and 34 μg/mL of catalase (bovine liver - lyophilized powder, Sigma-Aldrich, C40). Reducing agent was not included in the buffers.

**Video S1.** Cy5 FluoroCubes in STORM Buffer

Buffer: High-glucose DMEM containing 0.56 mg/mL glucose oxidase and 34 μg/mL catalase

Imaging conditions: 20 fps (50 ms exposure), RT, 640 nm illumination at ~0.56 $kW \cdot cm^{-2}$

**Video S2.** Cy5 FluoroCubes in Optimized Buffer

Buffer: High-glucose DMEM containing 0.56 mg/mL glucose oxidase, 34 μg/mL catalase, and 1× Trolox

Imaging conditions: 20 fps (50 ms exposure), RT, 640 nm illumination at ~0.56 $kW \cdot cm^{-2}$

**Video S3.** 40 nm Red-Orange (565/580) FluoSpheres in Optimized Buffer

Buffer: High-glucose DMEM containing 0.56 mg/mL glucose oxidase, 34 μg/mL catalase, and 1× Trolox

Imaging conditions: 20 fps (50 ms exposure), RT, 532nm illumination at ~0.22 $kW \cdot cm^{-2}$

**Derivation of the modified Kanade-Lucas-Tomasi (KLT) optical flow tracking algorithm**

**Optical flow tracking**

For the use of the standard KLT algorithm, one considers a pair of images at two time points denoted by $t$ and $t$+1. The whole image is partitioned into small interrogation windows, each containing $n$ pixels. Finding the motion of each individual interrogation window is formulated as an optimization problem. The pixel intensities $I(\mathbf{x}_i, t)$ at positions $\mathbf{x}_i$ are compared to pixel intensities $I(\mathbf{x}_i + \mathbf{u}, t+1)$, where all positions in the interrogation window are shifted by the same displacement vector $\mathbf{u}$ as

$$\arg\min_{\mathbf{u}} \sum_{i=1}^{n} (\|I(\mathbf{x}_i + \mathbf{u}, t+1) - I(\mathbf{x}_i, t)\|^2). \quad (1)$$

An expansion $I(\mathbf{x}_i + \mathbf{u}, t+1)$ is based on the assumption of small displacements $\mathbf{u}$ and yields in the first order the optical flow equations, which are given by

$$\underbrace{\begin{pmatrix} I_x(x_1, y_1) & I_y(x_1, y_1) \\ \cdots & \\ I_x(x_n, y_n) & I_y(x_n, y_n) \end{pmatrix}}_{\boldsymbol{A}} \begin{pmatrix} u_x \\ u_y \end{pmatrix} + \underbrace{\begin{pmatrix} I_t(x_1, y_1) \\ \cdots \\ I_t(x_n, y_n) \end{pmatrix}}_{-\mathbf{b}} = 0, \quad (2)$$

where $I_{\mathrm{x}}(x_i, y_i) = \frac{\partial I(\mathbf{x}_i, t)}{\partial x}$, $I_{\mathrm{y}}(x_i, y_i) = \frac{\partial I(\mathbf{x}_i, t)}{\partial y}$, $I_{\mathrm{t}} = I(\mathbf{x}_i, t+1) - I(\mathbf{x}_i, t)$. This linear system is typically overdetermined since the number of pixels $n$ is greater than the number of unknowns. Therefore, Equation 2 can be solved with a least-square approach as

$$(\boldsymbol{A}^{\mathrm{T}}\boldsymbol{A})\mathbf{u} - \boldsymbol{A}^{\mathrm{T}}\mathbf{b} = 0, \quad (3)$$

$$\mathbf{u} = (\boldsymbol{A}^{T}\boldsymbol{A})^{-1}\boldsymbol{A}^{\mathrm{T}}\mathbf{b}, \quad (4)$$

where

$$\boldsymbol{A}^{\mathrm{T}}\boldsymbol{A} = \begin{pmatrix} \sum_{i=1}^{n}\left(I_x(x_i, y_i)\right)^2 & \sum_{i=1}^{n}\left(I_x(x_i, y_i)I_y(x_i, y_i)\right) \\ \sum_{i=1}^{n}\left(I_x(x_i, y_i)I_y(x_i, y_i)\right) & \sum_{i=1}^{n}\left(I_x(x_i, y_i)\right)^2 \end{pmatrix}, \quad (5)$$

$$\boldsymbol{A}^{\mathrm{T}}\mathbf{b} = \begin{pmatrix} \sum_{i=1}^{n}\left(I_x(x_i, y_i)I_t(x_i, y_i)\right) \\ \sum_{i=1}^{n}\left(I_x(x_i, y_i)I_t(x_i, y_i)\right) \end{pmatrix}. \quad (6)$$

Equation 3 has a well-defined solution given by Equation 4 if $\boldsymbol{A}^{\mathrm{T}}\boldsymbol{A}$ is invertible, which requires $\mathrm{Tr}(\boldsymbol{A}^{\mathrm{T}}\boldsymbol{A}) \neq 0$. This condition is automatically satisfied by incorporating the minimum eigenvalue feature detection algorithm[2], which mandates that the eigenvalues of $\boldsymbol{A}^{\mathrm{T}}\boldsymbol{A}$ for a feature point are greater than a user-defined threshold. In practice, interrogation windows containing features comprising bright pixels can be used for tracking. The solution to Equation 3 is obtained via an iteration scheme proposed in Ref.[3].

**Tracking motion in dual-channel fluorescence images**

With our experimental setup, we record image pairs from both the beads and the FluoroCubes before and after removal of the adherent cell, producing two pairs of images that share the same displacement field. We test different approaches to track the displacements of the beads and FluoroCubes based on points detected independently from each channel. First, displacements in both channels are tracked individually using the conventional KLT algorithm, and the tracking results are combined to obtain a dense displacement field. Second, dual-channel images can be converted to single-channel images by weighted summation of the intensities. A simple choice is to assign the same weight to both channels and average the images as

$$I(x, y) = 0.5\, I^{(1)}(x, y) + 0.5\, I^{(2)}(x, y), \tag{7}$$

where the superscripts indicate the image channel. The resulting averaged image can used for motion tracking with the conventional KLT algorithm. Third, as an alternative approach for extracting information from two images channel, we modify the original KLT algorithm by extending the dimensions of the matrix $A$ given in Equation 2 as

$$\begin{pmatrix} I_x(x_1, y_1) & I_y(x_1, y_1) \\ \dots & \\ I_x(x_n, y_n) & I_y(x_n, y_n) \end{pmatrix} \rightarrow \begin{pmatrix} I_x^{(1)}(x_1, y_1) & I_y^{(1)}(x_1, y_1) \\ \dots & \\ I_x^{(1)}(x_n, y_n) & I_y^{(1)}(x_n, y_n) \\ I_x^{(2)}(x_1, y_1) & I_y^{(2)}(x_1, y_1) \\ \dots & \\ I_x^{(2)}(x_n, y_n) & I_y^{(2)}(x_n, y_n) \\ \dots & \end{pmatrix}, \tag{8}$$

where the superscripts on the matrix entries on the right side indicate the image channels. The KLT algorithm, used with the extended matrix, simultaneously tracks the displacements from both channels while imposing the constraint that the displacement field is the same in both channels. Additionally, this modification avoids cross-channel terms in the target function, which results in errors in densely labeled images, as discussed in the Results section. However, if one of the two image channels contains much more noise than the other, e.g., if some fluorescent signals occur in one frame but disappear in the next frame, the precision of the displacements estimated from both channels is degraded by the data from the channel with the larger noise. We prevent this problem by using an iterative tracking routine that imposes for every interrogation window in both channels a different weight that is based on previous tracking results. The choice of weighting factors is, in principle, flexible. Here, we use for the weighting factor the cross-correlation value associated with every tracked interrogation window with given displacement $\mathbf{u}$ for the two channels independently as

$$w^c = \frac{\sum_{i=1}^{n}(I^c(\mathbf{x}_i + \mathbf{u}, t+1) - \overline{I^c}(\mathbf{x}_i + \mathbf{u}, t+1))(I^c(\mathbf{x}_i, t) - \overline{I^c}(\mathbf{x}_i, t))}{\sqrt{\sum_{i=1}^{n}(I^c(\mathbf{x}_i + \mathbf{u}, t+1) - \overline{I^c}(\mathbf{x}_i + \mathbf{u}, t+1))^2 \sum_{i=1}^{n}(I^c(\mathbf{x}_i, t) - \overline{I^c}(\mathbf{x}_i, t))^2}} \tag{9}$$

where $c$ is the channel index, $i$ is the index denoting the pixels in a local window, and $\bar{I}$ is the mean intensity of the interrogation window. Then, the weighted matrix $\boldsymbol{A}$ becomes

$$\boldsymbol{A} = \begin{pmatrix} w^{(1)}I_x^{(1)}(x_1, y_1) & w^{(1)}I_y^{(1)}(x_1, y_1) \\ \cdots & \\ w^{(1)}I_x^{(1)}(x_n, y_n) & w^{(1)}I_y^{(1)}(x_n, y_n) \\ w^{(2)}I_x^{(2)}(x_1, y_1) & w^{(2)}I_y^{(2)}(x_1, y_1) \\ \cdots & \\ w^{(2)}I_x^{(2)}(x_n, y_n) & w^{(2)}I_y^{(2)}(x_n, y_n) \\ \cdots & \end{pmatrix}. \tag{10}$$

Ideally, a perfectly tracked point should have its maximum cross-correlation value at the center of the cross-correlation matrix in both channels. To use this condition for an assessment of the precision of each track, we define a threshold for the distance between the center of the cross-correlation matrix and the maximum of the cross-correlation values. A tracked point must be re-tracked if either the distance between the center and the maximum exceeds the threshold or if the absolute value of the maximum cross-correlation value is less than a second threshold. We repeat the tracking procedure with updated weights until a predefined termination criterion is reached, i.e., if all tracks are accepted according to the above criteria or if a maximum number of iterations is exceeded.
